# Proteins released into the plant apoplast by the obligate parasitic protist *Albugo* selectively repress phyllosphere-associated bacteria

**DOI:** 10.1101/2022.05.16.492175

**Authors:** Daniel Gómez-Pérez, Monja Schmid, Vasvi Chaudhry, Ana Velic, Boris Maček, Ariane Kemen, Eric Kemen

## Abstract

Biotic and abiotic interactions shape natural microbial communities. The mechanisms behind microbe-microbe interactions, particularly those protein-based, are not well understood. We hypothesize that secreted proteins are a powerful and highly specific toolset to shape and defend plant niches. We have studied *Albugo candida*, an obligate plant parasite from the protist Oomycota phylum, for its potential to modulate the growth of bacteria through secretion of proteins into the apoplast. Amplicon sequencing and network analysis of *Albugo*-infected and uninfected samples revealed an abundance of negative correlations between *Albugo* and other phyllosphere microbes. Analysis of the apoplastic proteome of *Albugo* colonized leaves combined with machine-learning predictors enabled the selection of candidates for heterologous expression and study of their inhibitory activity. We found that three of the candidate proteins show selective antimicrobial activity on gram-positive bacteria isolated from *Arabidopsis thaliana* and that these inhibited bacteria are important for the stability of the community structure. We could ascribe the antibacterial activity of the candidates to intrinsically disordered regions and positively correlate it with net charge. This is the first report of protist proteins with antimicrobial activity under apoplastic conditions that therefore are potential biocontrol tools for targeted manipulations of the microbiome.

## Introduction

The plant leaf is a highly competitive habitat for microbes due not only to limited resources but also to its instability as a result of rapidly changing conditions e.g., microbes triggering defense reactions or exploiting the habitat up to its destruction (Hassani et al., 2018). As a consequence, mechanisms that enable microbes to fight off opponents, by, e.g., outcompeting competitors for limiting resources or releasing antimicrobial compounds, are under strong selective pressure (Freilich et al., 2011). Identification and characterization of such mechanisms could lead to breakthroughs in therapeutics and disease control (Bollenbach, 2015). In particular, studies on stable interactions in natural microbial communities have historically been considered an important resource in the discovery of new antimicrobial compounds (Molloy and Hertweck, 2017). The best adapted microbes are obligate biotroph symbionts or pathogens that can only survive on a living host (Ruhe et al., 2016). They rely completely on intact plant niches where host and microbes are in stable equilibrium. In microbial community network analyses of the phyllosphere, hub microbes emerge as highly interconnected microbes that play a central role in the management of the microbial composition (Agler et al., 2016). The oomycete and obligate biotroph pathogen *Albugo* was shown to be such a microbe by reducing the growth of some microbes while increasing the growth of others and thereby significantly impacting the leaf microbial community (Agler et al., 2016). However, the mechanisms that underlie inhibition or promotion of co-occurring microbes remain largely unexplored. Therefore, *Albugo* infection and its effect on the microbiome represents an ideal model system to identify and study antimicrobial strategies in obligate pathogens that need to defend their niche to keep the host alive.

*Albugo* is the causal agent of white blister rust on Brassicaceae plants. Taxonomically, it belongs to the Oomycota, a heterogeneous group of protists comprising many highly adapted parasites of plants, animals, and humans. Following penetration into the plant host via the leaf surface, *Albugo* develops intercellular hyphae to colonize the plant extracellular space, known as apoplast (Berlin and Bowen, 1964). Herein, it is in contact with other endophytic microbes and competes for nutrient and habitat dominance. As an obligate biotroph, *Albugo* relies on the living plant for nutrients and structural support and hence, for overall survival. As a consequence, *Albugo* is incapable of growing independently of its host and reduction in its genome has led to the loss of all of its secondary and most of its primary metabolic pathways (Kemen et al., 2011). To shape its niche, *Albugo* secretes proteins into the plant cytoplasm and the apoplast. Some of these so-called effector proteins modulate host immune responses, but for many of them the function remains unknown (Furzer et al., 2022). As described only recently for a hemibiotrophic fungus, apoplastic secreted proteins can also act as potential microbiome control agents since they selectively modify the endophytic bacterial community and can therefore be considered effectors governing microbe-microbe interactions (Snelders et al., 2020).

Enrichment in long intrinsically disordered regions (IDRs) is a common feature of the secretome of plant pathogens (Marín et al., 2013). IDRs are protein domains that, in general terms, lack a stable folding conformation (Oldfield et al., 2019). This is due to their sequence, which is biased towards certain disorder-promoting amino acids and often shows hydrophilic tendencies (Dubreuil et al., 2019). The function of IDRs in proteins can vary considerably depending on the environmental conditions, owing to their inherent structural pliability. In plant pathogenic eukaryotes, IDRs have been proposed to be relevant for extracellular effector protein delivery into host cells or the apoplast (Liu et al., 2019). Effectors need to be flexible enough to evade host recognition and require a certain plasticity to bind host targets even with slight variations (Marín et al., 2013). Recently, IDRs have been found to be responsible for antimicrobial activity, particularly in peptides with a positive net charge (Latendorf et al., 2019). These cationic intrinsically disordered antimicrobial peptides (CIDAMPs) could be a novel source of highly specific antimicrobials, especially those from obligate biotrophs, as they do not harm the host but specifically shape the niche for the needs of the pathogen.

Here, we explore the antimicrobial activity associated with IDRs of apoplastic proteins from *Albugo candida*, and report for the first time an example of a protist and obligate biotrophic pathogen as a potential source for highly specific antimicrobials.

## Results

### *Albugo* is highly intercorrelated

We set out to predict robust and ecologically relevant correlations between *Albugo* and other co-occurring microbes, as well as assess the significance of the number of these correlations in the context of the phyllosphere. With these aims, we applied the software FlashWeave to infer direct interactions between the operational taxonomic units (OTUs) of a large amplicon sequencing dataset of wild *A. thaliana* from six consecutive years in six locations (735 samples containing a total of 11,172 OTUs). In the resulting correlation network, consisting of 123,316 edges and 11,150 OTUs, we found the *Albugo* sp. OTU to be in the upper top 0.85 quantile of the total interactions with a degree of 33 including 21 positive interactions (Figure S1). Furthermore, this OTU was in the top 20 when ranked by negative interactions with a total of 12, including connections to bacteria, fungi and one other eukaryote (Figure 1a, Figures S2 and S3). The negatively correlated bacteria included mostly gram-negative strains of which the most abundant phylum was the Proteobacteria with four members. The negatively correlated eukaryotes included an ascomycete fungus and two green algae (Figure 1b). In summary, our analysis indicates that the protist pathogen *Albugo* is highly intercorrelated in the phyllosphere, particularly through negative correlations when compared to other plant microbes.

**Figure 1.**
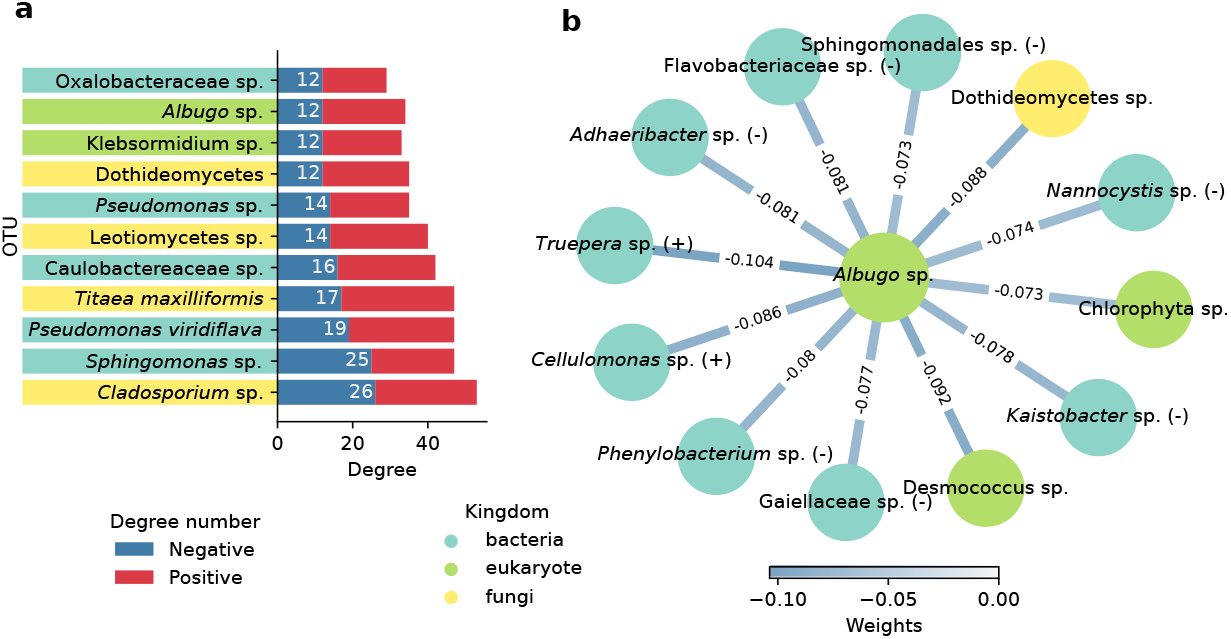
Network of operational taxonomic unit (OTU) interactions in the phyllosphere of *Arabidopsis thaliana*. **(a)** Degree distribution of OTUs in the interaction network, showing nodes with 12 or more negative interactions and more than 100,000 total reads in the dataset, where *Albugo* places tenth. (**b**) Inferred negative interactions for the *Albugo* sp. OTU. Color of the edges and nodes represent the strength of the correlation and the phylogenetic kingdom to which the node belongs, respectively. Unless explicitly stated, taxonomy at the species level could not be resolved with confidence (bootstrap *<* 90). Gram-stain of bacterial nodes is displayed in parentheses.

### Antimicrobial proteins are enriched in the apoplast

To identify potential, protein-based causal agents of such negative correlations resulting in a reduction of microbial diversity as previously described by (Agler et al., 2016), we studied *Albugo* secreted proteins in the plant apoplast (Figure 2a). Through proteomic analysis, we identified 563 proteins from *A. candida* in the apoplast of infected *A. thaliana* leaves with at least one peptide at an Andromeda score higher than ten, representing 4.2% of the total predicted proteome of *A. candida* (Figure 2b). Among these, 70 proteins (12.4%) carried a putative secretion signal and 12 had a mitochondrial inner membrane localization according to the GO assignment compared to 28 in the predicted proteome of *A. candida*, suggesting minor contamination from broken hyphae. After performing annotation of biological processes through GO terms, we found that 429 out of the 563 proteins (76.2%) resulted in significant hits (Figure S4). For these, we studied the enrichment of biological functions compared to the predicted intracellular proteome of *A. candida*. Carbohydrate, amino acid and nucleic acid, catabolism and biosynthesis featured prominently in the enriched terms (Fisher’s exact test Holm-corrected *p* value *<* 0.001; Figure 2c). Additionally, we employed a de novo antimicrobial activity prediction approach in order to find antimicrobial proteins with no known conserved regions. We found two of the tools (AmpGram and amPEPpy) to be biased towards a longer or shorter protein length in our dataset, therefore we adjusted the weight based on the R-squared value of the correlation (Figure S5). Following the weighted score, a total of 154 apoplastic proteins were found to be positive for antimicrobial activity (27.3%, compared to 26.3% in the entire *A. candida* proteome). This corresponded to a significant enrichment for the presence of predicted antimicrobial proteins in the apoplastic proteome (Mann-Whitney U test, *p* value = 0.013; Figure 2d).

**Figure 2.**
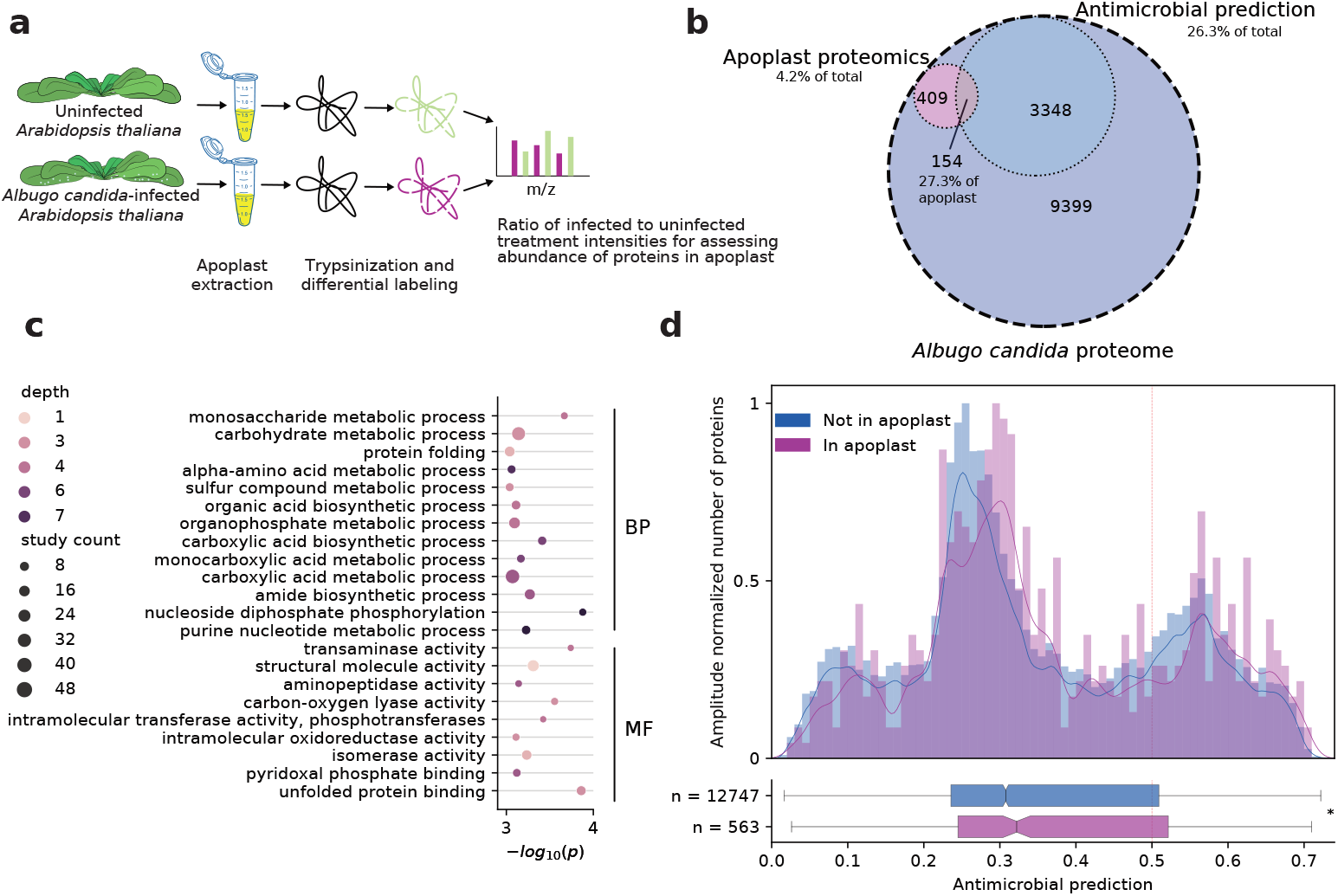
Proteomics of *Albugo candida* apoplastic proteins from infected *Arabidopsis thaliana* leaves. (**a**) Workflow of the proteomics analyses on infected (10 days post infection) and uninfected *Arabidopsis thaliana*. (**b**) Venn diagram of *A. candida* proteome. A total of 563 proteins were identified in the apoplast, of which 154 have a predicted antimicrobial function. (**c**) Enrichment of gene ontology biological processes related to biological processes (BP) and molecular functions (MF) in apoplastic proteins compared to background proteome in *A. candida* with a significance of *p* value *<* 0.001. (**d**) Normalized histogram of antimicrobial predictions in proteins found or not in the apoplastic proteomics for *A. candida* with kernel density estimate represented as a line. At the bottom, a box plot represents the distribution of proteins with antimicrobial prediction within the apoplastic and non-apoplastic subset (Mann-Whitney U test, *p* value = 0.015). A dashed red line indicates the cutoff of 0.5.

To corroborate and study the potential antimicrobial properties of these proteins in more detail, we selected candidates for overexpression in a heterologous system. We considered the following properties when choosing candidate proteins: 1. Positive prediction for antimicrobial activity (cutoff higher than 0.5 for consensus prediction), 2. High abundance in the apoplast as measured by the relative peptide intensity compared to that of the uninfected treatment in the proteomics (above median of the normalized distribution), and 3. Short sequence length to match the upper bound of known effector proteins (less than 600 amino acids). As negative controls, we selected proteins with a lower antimicrobial prediction that had a comparable size and abundance to the antimicrobial candidates. We were able to amplify representative candidates with a positive (C06 and C14) and a negative (C05 and C15) prediction for antimicrobial activity using as template a cDNA library of *A. candida*-infected *A. thaliana* (Table S1). Of note, the predicted peptide signal for classical secretion of C14 and C15 was removed during cloning. The candidates were subsequently heterologously expressed in *Escherichia coli* to test in vitro the antimicrobial activity of the corresponding recombinant proteins as described below.

### Heterologously expressed candidates show antimicrobial activity

During overexpression of the candidates in the *E. coli* system, we observed accumulation of C05 and C14 in inclusion bodies after IPTG induction under standard conditions. By systematically testing different expression settings, including lower temperature (15 °C to 37 °C), lower inducing concentration (0.1 mm to 1 mm IPTG) and longer induction time (4 h to 48 h), we could natively extract C06 and C15 as soluble proteins. Extraction under denaturing conditions using urea was successful for all proteins, regardless of whether they were synthesized into inclusion bodies or not (Figure S6). Therefore, we used a denaturing extraction protocol as the standard purification method for comparison of all the expressed candidates (Figure S7). After purification and concentration of the candidates in a testing buffer resembling the apoplastic pH conditions (BisTris-based buffer at pH 5.9), we performed an antimicrobial screen on a selection of 24 strains from an in-house microbial strain collection of plant-isolated bacteria that were cultured under standard conditions from *A. thaliana* samples (Tables S3). These strains were selected based on a pre-screen which revealed gram-positive bacteria to be more sensitive to the protein treatment than gram-negative. The microbial collection also includes strains that were detected as core members in the *A. thaliana* phyllosphere microbiome, meaning they are consistently present in natural communities and hence are likely to inhabit the plant before or during an *Albugo* infection (Almario et al., 2022). Overall, we observed a variable effect on the growth of bacteria, with species within the same genus showing different responses. We found selective antimicrobial activity against five strains, including two *Exiguobacterium* (I10, I11), a *Curtobacterium* (I06), an *Aeromicrobium* (I01) and a *Microbacterium* (I20). To a lesser extent, we observed antimicrobial activity on four other strains, a *Sanguibacter* (I30), a *Plantibacter* (I24) and two *Microbacterium* (I17, I21; Figure 3b, S8 and S9).

**Figure 3.**
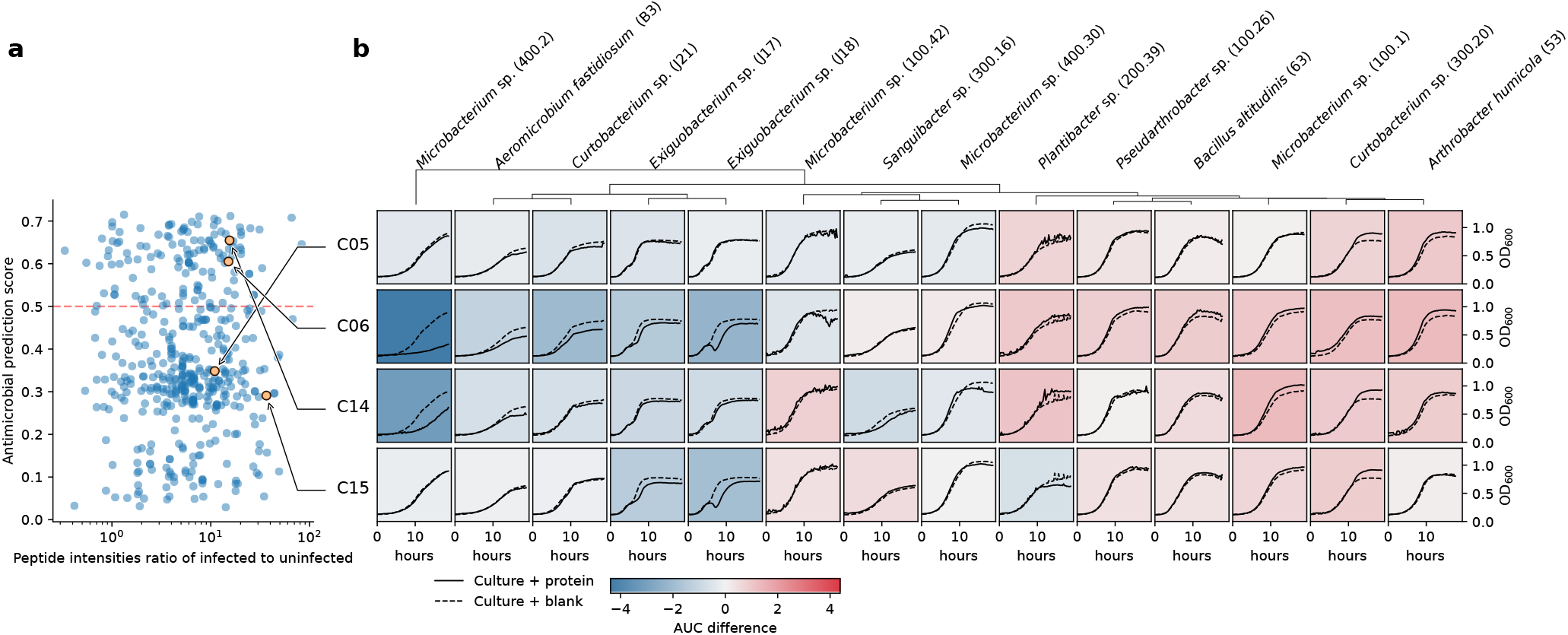
Antibacterial prediction and activity of apoplastic candidate proteins from *Albugo candida*. **(a)** Antimicrobial prediction and relative intensity of the proteins found in the apoplast of *Arabidopsis thaliana*, highlighting the selected candidates. (**b**) Growth curves for candidates C05, C06, C14 and C15 at a concentration per well of 0.75 μM compared to blank (dashed lines) during 19 h of growth. Background color represents inhibition (blue) or promotion (red) of growth based on the difference in the area under the curves. 95% confidence intervals for these tests are shown in Figure S9.

Candidate C06 showed the highest inhibitory activity, while C14 and C15 showed minor inhibition at equal molarity (0.75 μM). Based on the in silico prediction, we expected C06 and C14 but not C15 to display antimicrobial activity (Figure 3a). C05, instead and consistent with the antimicrobial prediction, had the least antimicrobial effect of all. We additionally found that all proteins displayed a variable growth promoting effect towards most other gram-positive strains: a *Curtobacterium* (I07), a *Pseudarthrobacter* (I25), a *Microbacterium* (I18), a Streptomyces (I36), an *Exiguobacterium* (I09), two *Bacillus* (I04, I05) and two *Arthrobacter* (I02, I03). The five tested gram-negative strains remained mostly unaffected by incubation with the protein candidates during growth (Figure S8). We also tested the antimicrobial activity at the higher pH of 7.2 to see whether apoplastic conditions were necessary for the inhibitory effect to take place or if cytoplasmic conditions are more likely to enhance antimicrobial function. We found a significant loss of antimicrobial activity for C06 at the pH of 7.2 (Figure S10). Thus, in summary, we found a high correlation of the consensus prediction method with the experimental antimicrobial results exclusively at a pH reflecting apoplastic conditions.

Computational analysis of all predicted antimicrobial proteins that had been identified by proteomics in the apoplast revealed a significant enrichment for IDRs compared to non-apoplastic proteins (Fisher’s exact one-tailed test, *p* value = 0.002). This, together with the highly enriched term for unfolded protein binding (Figure 2c), suggests the importance of disordered regions in the secreted proteome. Consistent with this, the candidates that showed antimicrobial activity had long predicted IDRs, notably in the C-terminus with a positive net charge (Figure 4a). C15, which was not predicted as an antimicrobial, did also present putative IDRs but they were slightly shorter in extension at the N-terminus (maximum stretch of 23 amino acids, vs 129 for C06 and 30 for C14). Additionally, C06 had a compositional bias for alanine and glutamine, and C14 and C15 had a bias towards increased serine (Table S5), all of which are disorder-associated amino acids (Uversky, 2013). To test whether the positively charged IDRs were responsible for the antimicrobial activity, we separately cloned and purified the C-terminal regions of C06 and C14 (165 and 129 amino acids long, respectively), which were both predicted to contain IDRs and display high positive net charge (Table S2), as well as the remaining N-terminal region of each protein, which was for the most part not predicted to be disordered.

**Figure 4.**
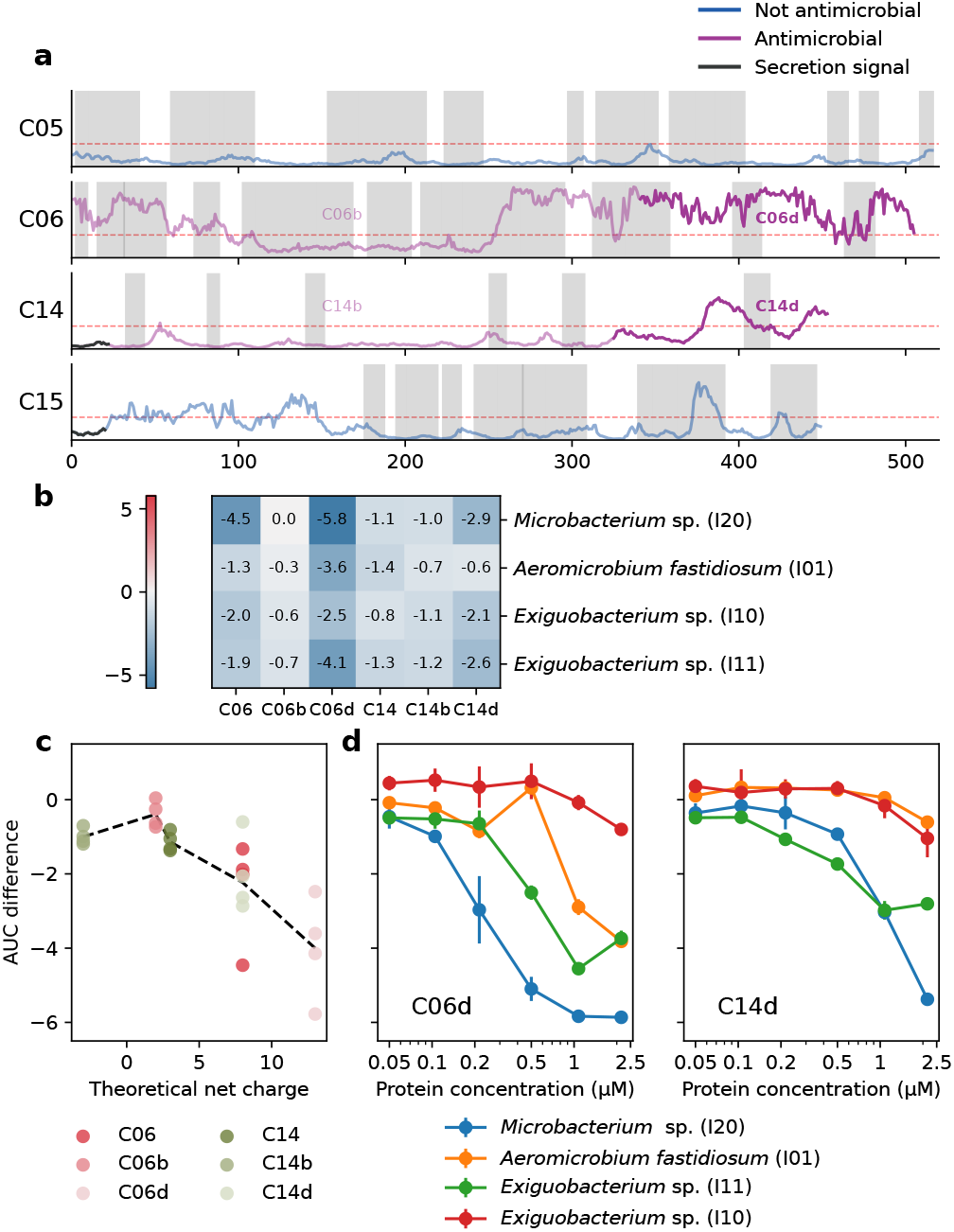
Antimicrobial activity of protein candidates from *Albugo candida*, C06, C14 and their domains. **(a)** Individual candidate predictions of disorder per amino acid, emphasizing the annotated domains used for antimicrobial testing in bold (domain d) or faded color (domain b). The threshold for positive disorder (0.3) is marked in red. The secretion signal of C14 and C15 is depicted in gray. Shaded regions represent the peptide coverage as found in the proteomics analyses. (**b**) Comparison of the antimicrobial activity of the different domains of C06 and C14 at a concentration of 1 μM shown as the differences in the area under the curve to blank during 19 h of growth. The values represent the mean of at least three biological replicates. Full curves are found in Figure S11. (**c**) Antimicrobial activity as measured by the change in the area under the curve (AUC) correlated to predicted net charge for all domain tests with LOWESS regression showing a downwards trend line. (**d**) Concentration dependent inhibitory activity of C06 and C14 C-terminal disordered domains towards four sensitive strains. Vertical bars represent standard deviation. Full curves are found in Figure S12.

We found the strongest antimicrobial activity for both disordered C-terminal domains of C06 and C14 (C06d and C14d) when compared to the whole protein and the N-terminus domains (Figure 4b and Figure S11). We observed a correlation between the predicted net charge of the peptides and proteins of C06 and C14 and their antimicrobial effect: the higher the net charge, the higher the antimicrobial activity (Figure 4c and Figure S11). Furthermore, for C06d and C14d we observed a concentration dependent effect against *Exiguobacterium* strains I10 and I11, *Aeromicrobium fastiodiosum* strain I01 and *Microbacterium* strain I20, with C06d reaching a complete inhibitory activity against *Aeromicrobium* and *Microbacterium* growth at 2.15 μm (Figure 4d and Figure S12). In the proteomics dataset, we found not only coverage for a large part of the sequence of both positive candidates (Figure 4a) but the identified peptides also indicated the presence of the disordered domains with antimicrobial activity in the apoplast, particularly for C06d (Table S6). To support the hypothesis that the proteins might get cleaved allowing the release of the IDR peptides in the apoplast we used the proteomics dataset to study potential changes in *A. thaliana* protease activity upon infection. We found evidence of an upregulation of proteases in the apoplast, particularly subtilisin-like serine proteases, in the presence of *A. candida* infection (Table S7). Hence, we conclude that the observed antimicrobial function of C06 and C14 might be traced back to the IDRs in the C-terminus, which is in line with reports from human proteins that contain IDRs, the CIDAMPs.

To see whether C06d, as the strongest inhibitor, could in principle be used as an effective microbial control agent, we conducted antimicrobial assays with three grampositive strains isolated from *A. thaliana* that were closely related to known plant pathogens (Table S6). These included *Rhodococcus fascians* (Dhaouadi et al., 2020; Hjerde et al., 2013), *Clavibacter michiganensis* subsp. *tessellarius* (Carlson and Vidaver, 1982; Li and Yuan, 2017) and subsp. *capsici* (Oh et al., 2016). As in the previous tests, we found the inhibition to be strain specific. While *R. fascians* (I45, I46) and *C. michiganensis* subsp. *tessellarius* (I43, I44) were unaffected by protein addition, all tested isolates from *C. michiganensis* subsp. *capsici* (I37-I42) showed a variable degree of growth inhibition (Figure S13). Therefore, due to its ability to inhibit phytopathogenic *Clavibacter*, C06d would present a viable candidate for further investigation regarding its role as a potential pathogen-control agent.

To examine whether the peptides showed inhibitory activity within a microbial community context, we conducted SynCom experiments in culture media. A SynCom composed of *Aeromicrobium* (I01), which was inhibited in the previous tests, and four additional strains (I03, I04, I13, I28), all unaffected in their growth by protein addition, was constructed (Figure S14). We added a rifampicin-resistant pathogen *Pseudomonas syringae* to use as read-out. We found that the growth of Pst was increased when *Aeromicrobium* was excluded from the SynCom, as compared to the whole five-strain SynCom, when grown with testing buffer (Figure 5a and S15). To see if C06d could effectively inhibit *Aeromicrobium* in a multi-strain community, we added the protein solution. In this setting, the growth of Pst was comparable to the *Aeromicrobium* dropout treatment, indicating inhibition by C06d. As a control, we also tested the SynCom with C06b, which showed no inhibitory activity (Figure S14). The Pst load, as measured by the CFU count per mL was similar to the five-strain SynCom treatment in buffer (Figure Figure 5a and S15). Together, these results indicate that the inhibitory properties of C06d are persistent in a multi-strain bacterial setting.

**Figure 5.**
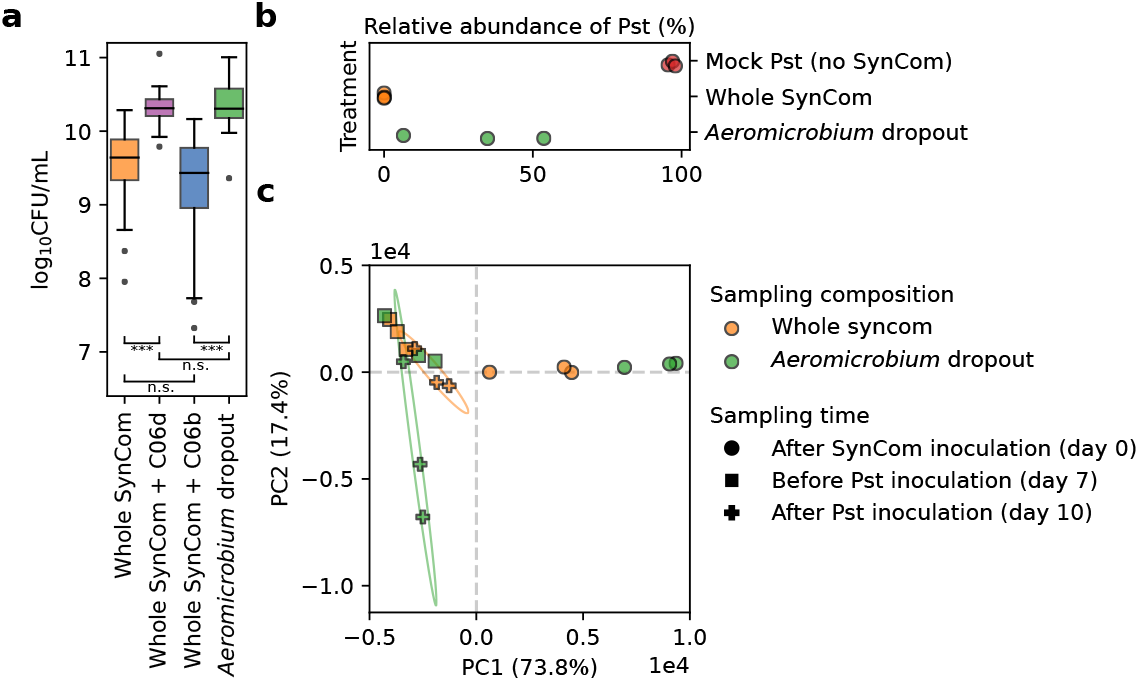
Synthetic community (SynCom) experiments in liquid culture and in planta suggest relevance of *Aeromicrobium fastidiosum* IO1. **(a)** C06d (purple) and C06b (blue) 1 μM protein treatments on a five-strain bacterial community in liquid culture result in variable community disruption as measured by the bacterial load of *Pseudomonas syringae* pv. tomato DC3000 (Pst) in CFUmL^−1^. Whole SynCom with buffer control (BisTris, pH 5.9) is shown in orange, *Aeromicrobium fastidiosum* (I01) dropout community in green. Significance tests show a difference between whole SynCom treatment and C06d treatment, as well as C06b treatment and *Aeromicrobium* dropout (Mann-Whitney U; n.s.: not significant, ***: *p* value < 0.001). Detailed counts per replicate can be found in Figure S15. **(b)** Protective effect of a 15-strain SynCom with or without *Aeromicrobium fastidiosum* (I01) as measured in planta by the relative abundances of Pst three days post infection. Mock treatment represents plants not treated with the SynCom. (**c**) Principal coordinate analysis of the amplicon sequencing variant (ASV) frequencies in the SynCom dropout experiment on *Arabidopsis thaliana*. Calculated based on the Euclidean distances of bacterial 16S rRNA ASVs abundances. Ellipses drawn for the final timepoints based on two standard deviations show a large variation for the *Aeromicrobium* dropout but not for the whole SynCom after Pst disruption.

### Inhibited microbes are important for community stability

To test for the relevance of the inhibited microbes in a natural ecological context, we assessed their necessity for community stability in planta. We performed dropout experiments on a SynCom in *A. thaliana* consisting of 12 bacterial strains and three yeasts, all identified as core microbes, challenged with a model plant-pathogenic strain of *Pseudomonas syringae*. If in the SynCom, *Aeromicrobium fastidiosum* (I01) was not included, we observed a variable decrease in the resistance of the plant to infection (Figure 5b). We also observed an overall increase in the sample-to-sample variation of the bacterial abundances compared to the whole SynCom treatment, which were consistent for the three replicates. This was generalizable to the entire bacterial composition of the samples as measured by the lack of clustering in the principal coordinate analysis for the *Aeromicrobium* dropout (Figure 5c). Additionally, through pairwise interaction assays with SynCom members, this *Aeromicrobium* strain was shown to inhibit other bacterial strains. This highlighted *Aeromicrobium*’s potential in stabilizing the microbiome by controlling the abundance of other core microbes in the phyllosphere of *A. thaliana* (Figure S16). Furthermicrobial lawn experiments of *Aeromicrobium* showed its resilience against other SynCom members in one-to-one interactions (Figure S17).

## Discussion

The plant apoplast is a challenging habitat for the survival of microbes. On the one side, plant defenses, including oxidative bursts and proteases, and on the other, microbes defending their niche make it a difficult environment to conquer (Jashni et al., 2015; Wang et al., 2020). *Albugo* is a filamentous pathogen that has developed a niche-forming strategy, whereby it modifies the pre-existing plant microbiome to better fit its needs. Some mechanistic explanations for this phenomenon have been attributed to *Albugo*’s effect on the immune system of the plant and its indirect repercussions on other microbes (Cooper et al., 2008; Ruhe et al., 2016). Specifically, by reducing the plant’s defense recognition systems, the pathogen allows the thriving of microbes that otherwise would not, or vice versa. Here, we provide evidence suggesting that *Albugo* directly contributes to shape the plant-associated microbial communities through the release of proteins and peptides with antimicrobial activity into the apoplast.

The OTU interaction analysis of the *A. thaliana* microbiome dataset showed a significant number of positive and negative interactions of *Albugo* with specific community members, supporting previous findings (Agler et al., 2016). The adjacent positive interactions imply a promotion of these organisms as potentially beneficial for *Albugo*. Within the network of positive interactions we found a correlation between *Albugo* and the bacterial genus *Variovorax*, which, as a common plant growth-promoting rhizobacterium, could promote survival of the host plant during infection, thus indirectly promoting *Albugo* (Chen et al., 2013; Finkel et al., 2020). In contrast, the numerous direct negative interactions can be explained by *Albugo*’s need to outcompete microbes that may be detrimental to its own survival (Figure 1b). Although the network is undirected and negative interactions could mean a repression by *Albugo* or the opposite, reduced microbial diversity in the phyllosphere upon *Albugo* infection points to a majority of negative connections being outgoing rather than ingoing (Agler et al., 2016). In agreement with this, *Albugo*, as an obligate biotroph, also has an interest in controlling the growth of microbes threatening survival of the host. By reducing the plant’s defenses through secretion of effectors, *Albugo* may leave the plant vulnerable to certain pathogens, which may have resulted in an adaptive antimicrobial response. For instance, *Cellulomonas*, which is shown in the network to be negatively correlated with *Albugo*, is a bacterial genus known for its plant cell wall degrading capabilities (Aydogan et al., 2018; Carlos et al., 2018). To keep the plant alive while suppressing its defense in the presence of numerous facultative pathogens is a general problem biotroph symbionts and pathogens face. In any case, they need to be able to control all those microbes, making them a currently unexploited resource for novel mechanisms of antimicrobial strategies.

One strategy to suppress competitors is the release of secondary metabolites. The genome of *Albugo*, however, does not contain key secondary metabolites potentially responsible for the observed broad spectrum of negative correlations with other microbes (Kemen et al., 2011). In fungi, potential protein effectors for microbe-microbe interactions were identified that have direct or indirect inhibitory effects on competitors (Eitzen et al., 2021; Snelders et al., 2020). For example, a fungal yeast has been shown to secrete a glycoside hydrolase responsible for inhibition of *Albugo* (Eitzen et al., 2021). Functional annotation of about 80% of the *A. candida*’s apoplastic proteins revealed a significant enrichment for metabolism-related processes. This might be explained by the sampling time point, that is, ten days after infection, when a high metabolic turn-over is required by the oomycete due to the active growth of hyphae. Regarding glycosyl hydrolases with potential antimicrobial function, *Albugo* has a significantly lower number compared to hemibiotrophic and necrotrophic oomycetes and only a small fraction of those could be identified in the predicted secretome and in the apoplastic secretome (Kemen et al., 2011). It has been shown that this is a common feature of obligate biotrophic pathogens, as lytic enzymes are potentially problematic since they may result in small molecular products that are recognized and trigger defense, eventually destroying the habitat (Zhang and Zhou, 2010). Thus, selection may have resulted in adaptation of proteins for this purpose. In this vein, C06 was unique in showing hints of positive selection as analyzed in a previous study, which may indicate recent selective adaptation (Gómez-Pérez and Kemen, 2021). We therefore hypothesized antimicrobial proteins or peptides as a mechanism to defend such a fragile niche within the leaf as these might be able to evade plant recognition but nevertheless restrict the growth of other microbes.

Machine learning has been used successfully to predict de novo antimicrobial activity during the screening and rational design of novel anti-infectives (Plisson et al., 2020). With the high-throughput approach applied in this study, we found a large percentage of proteins from *Albugo*’s predicted proteome to display putative antimicrobial activity, particularly those in the apoplastic dataset. Interestingly, all predicted antimicrobial proteins that were found in the apoplast displayed a significant enrichment in IDRs (Figure S18). Additionally, in the GO enrichment of molecular functions we found the term unfolded protein binding (Figure 2c). We hypothesize that apoplastic localization together with the presence of long IDRs and a positive net charge are good predictors for antimicrobial proteins and peptides. In our experiments, this was illustrated by C15, which despite the negative antimicrobial prediction, had IDRs and specific antimicrobial activity.

We found that C06 showed antimicrobial activity at pH 5.9 but not at pH 7.2, where it is expected to have a much more negative net charge (Table S2). In this manner, the pH could act as an external trigger of the antimicrobial effect exclusively in the more acidic conditions of the apoplast, thus preventing unwanted effects in the cytoplasm of the hyphae. Analogously, such a targeted mechanism has been reported for human antimicrobial peptides (AMPs), which are exclusively activated when they reach their site of action. This corresponds to the surface of the skin, which is an acidic environment with a comparable pH to that of the apoplast (Malik et al., 2016). Once secreted into the apoplast, *Albugo* proteins may be cleaved by proteases of plant or microbial origin, allowing for the release of AMPs and ensuring the full antimicrobial activity is only reached in proximity to the intended bacterial targets. IDRs may facilitate this process through the prevention of a globular conformation that restricts the access of proteases. Consistent with this hypothesis, we found evidence for an enrichment of *A. thaliana* proteases in the apoplast of *A. candida* infected plants (Table S7).

The antimicrobial effect of the proteins was much stronger when the IDR-rich domains were tested (C6d and C14d), while the less disordered domains (C06b and C14b) had the lowest activity at the same molarity (Figure 4b). The antimicrobial activity of cationic peptides has been ascribed to their interaction with cell envelopes which is facilitated by positively charged residues such as arginine, lysine and histidine (Cutrona et al., 2015). Of note, candidate C06 had a significant compositional bias for the presence of histidine (Table S5). As it was the case for the inhibited microbes in this study, particularly susceptible to cationic peptides are gram-positive bacteria due to the generally larger presence of negatively charged phosphatidylglycerol in their membrane compared to gramnegative. Thus, the variable antimicrobial effects of cationic peptides can be explained by the charge of the target membrane (Malanovic and Lohner, 2016). This mechanism could translate into contexts other than the *Albugo*-*Arabidopsis* pathosystem, allowing for the possibility of employing these peptides as inhibitors against strains of interest, e.g., phytopathogens like *C. michiganensis* subsp. *capsici*.

When looking at the bacteria inhibited by the *Albugo* protein candidates, we found them to have a large community-shaping potential. One of the highly inhibited strains, *Aeromicrobium fastidiosum* (I01), was responsible for a large part of the community stability as measured by the relative bacterial abundances after dropout in a synthetic community of *A. thaliana* following disruption with the bacterial phytopathogen Pst (Figure 5b and c). In line with observations from Agler et al. (2016), *Albugo* infection results in a reduced alpha diversity in the phyllosphere community. Specifically releasing protein effectors that target bacteria with an influence over a large part of the community may thus be a cost-effective way for *Albugo* to amplify its phyllosphere-shaping effect while having a reduced metabolic capability. The ability of the antimicrobial peptide C06d to inhibit bacteria in a complex in vitro community, further adds to this hypothesis (Figure 5a).

In summary, we have found three apoplastic proteins from the plant protist pathogen *A. candida* to be antimicrobial on several plant-isolated strains of gram-positive bacteria. These proteins were selected after apoplast proteomic analysis of leaf samples and in silico classification of the proteins in search for candidates with antimicrobial potential. Although their specific mechanism of action remains to be elucidated, we found a correlation of the antibacterial activity with the positive net charge in the IDRs of the C-terminal domains of two of these proteins. Given the large diversity of yet unexplored obligate biotrophs, this study opens the way for novel sources for the discovery of peptide-based antibiotics.

## Methods

### Interaction network inference

We inferred interaction correlations between OTUs on an amplicon sequencing dataset of *Arabidopsis thaliana*’s phyllosphere sampled twice a year from 2014 to 2019 in several sites around the area of Tübingen, Germany. Microbiome samples were isolated from the endo- or epiphytic compartment separately. The dataset combined analyses of 16S and 18S ribosomal RNA phylogenetic marker genes as well as of the ITS regions. We analyzed the raw OTU tables (Mahmoudi et. al, 2022, in preparation) that were constructed after a 97% similarity OTU clustering, using the software FlashWeave with a 0.05 significance cutoff, controlling for confounding meta variables including season, year, sampling site and phyllosphere compartment (Tackmann et al., 2019). We considered OTUs with a taxonomy assignment bootstrap below 90 as unclassified for that taxonomy level.

### Plant growth and infection

We infected *A. thaliana* plants of ecotype Ws-0, which had been grown under short day conditions for five weeks (8 h of light at 21 °C and 16 h of darkness at 16 °C), with *A. candida* strain Nc2 by spore suspension spray-inoculation. To obtain the latter, we submerged leaves with visibly sporulating *A. candida* pustules (at least 12 days post infection, dpi) in sterile ultrapure water for one hour and filtered the solution through miracloth (pore size of 25 μm). Following a cold treatment at 8 °C overnight and condensing humidity, we kept the inoculated plants under long day conditions (12 h of light and 12 h of darkness).

### Sample preparation for mass spectrometry analysis

We extracted apoplast samples from *A. candida*-infected and uninfected *A. thaliana* leaves at 10 dpi using vacuum pump infiltration at 100 Pa with 180 mm MES buffer (pH 5.5). We collected the infiltrate through centrifugation in a swinging-bucket centrifuge at 1000 RCF for 20 min. We performed acetone-methanol precipitation overnight at −20 °C. We resuspended the protein pellets in a denaturation buffer (6 m urea, 2 m thiourea, 10 mm Tris, pH 8), and determined their concentration by Bradford assay. Disulfide bonds were reduced with 10 mm DTT before alkylation with 55 mm iodoacetamide, for one hour each. We used trypsin (Promega Corporation) for digestion (Zittlau et al., 2021), and purified the peptides on Sep-Pak C18 Cartridges (Waters).

### High pH fractionation and dimethyl labeling

The Pierce High pH Reversed-Phase Peptide Fractionation Kit (kit no.: 84868; Thermo Fisher Scientific) was used to fractionate 150 μg of triple dimethyl-labeled and mixed proteome. Spin columns were conditioned twice with ACN and 0.1% TFA according to the vendor. We loaded the acidified peptides on columns and washed them once with water prior to elution. Peptides were eluted stepwise with 5%, 7.5%, 10%, 12.5%, 13.3%, 15%, 17.5%, 20% and final 50% ACN/ammonia. We acidified the fractions to pH 2 to 3 and desalted them on C18 StageTips before starting LC-MS/MS measurement. In total, we collected nine fractions per replicate. As previously described, we labeled the peptides with dimethyl on Sep-Pak C18 Cartridges (Boersema et al., 2009). We mixed triple dimethyl sets at an equal peptide ratio of 1:1:1. We inspected label efficiency and mixing in separate LC–MS/MS runs.

### LC-MS analysis

We analyzed the samples on a Q Exactive HF mass spectrometer (Thermo Fisher Scientific), and on an Exploris mass spectrometer (Thermo Fisher Scientific). An online-coupled Easy-nLC 1200 UHPLC (Thermo Fisher Scientific) was used to separate the peptides on a 20 cm analytical column, 75 μm ID PicoTip fused silica emitter (New Objective). This column was in-house packed with ReproSil-Pur C18-AQ 1.9 μm resin (Dr Maisch GmbH Ltd). We generated a gradient by solvent A (0.1% FA) and solvent B (0.1% FA in 80% ACN), at 40 °C and a 200 nL min^−1^ flow rate. We eluted the peptide fractions using a 90-min segmented linear gradient. We electro-sprayed and analyzed eluted peptides in a positive ion, data-dependent acquisition mode. We selected the top 12 most intense peptides. Full mass spectrometry was acquired in a scan range of 300 to 1750 m/z at a resolution of 60,000.

### MS data analysis and statistical analysis

We processed raw data files with the MaxQuant software suite version 1.6.7.0 (Cox and Mann, 2008). We searched the MS/MS data against the UniProt *A. thaliana* database (18,218 entries), the *A. candida* strain Nc2 predicted proteome, assembly accession: GCA_001078535.1 (Links et al., 2011) and commonly observed contaminants. We kept the search parameters to default values except for the dimethylation for light (28.03 Da), intermediate (32.06 Da), and heavy (36.08 Da) labels on lysine residues and peptide N-termini. We set oxidation of methionine and protein N-terminal acetylation as variable modifications and allowed carbamidomethylation of cysteine residues as fixed modification. All searches were performed in trypsin/P-specific digestion mode. Maximum of two missed cleavages were allowed. We considered an *Albugo* protein as present in the apoplast with a high confidence when it had at least one predicted peptide match with an Andromeda score higher than ten (Cox et al., 2011). We assessed the relative abundance of proteins in the apoplast by the normalized ratio of the peptide intensities between infected and non-infected treatments.

### Protein annotation and prediction

For the functional annotation, we used InterProScan version 5 (Jones et al., 2014). We analyzed the assigned gene ontology (GO) terms using the GOATOOLS software (Klopfenstein et al., 2018). For the antimicrobial assessment, we ran the predicted proteome of Nc2 through an antimicrobial prediction pipeline (https://github.com/danielzmbp/appred) consisting of several tools: Antimicrobial Peptide Scanner vr. 2, AmpGram and amPEPpy (Burdukiewicz et al., 2020; Lawrence et al., 2020; Veltri et al., 2018). These machine learning-based models attempt de novo prediction of antimicrobial activity not by similarity but by the compound features of the amino acid sequence. We added a weight to each method to account for their correlation to protein size, corresponding to one minus the absolute of the R-squared value when the leastsquares linear regression was significant. We considered a prediction as positive for antimicrobial activity when the weighted average of the three methods was higher than 0.5. For the prediction of disordered regions, we used the tool flDPnn with default settings and cutoff of 0.3 (Hu et al., 2021). We considered a protein to contain IDRs when it presented at least 15 predicted disordered residues in a consecutive order. We calculated the molecular properties of the candidates, including theoretical isoelectric point and molecular weight using Expasy’s ProtParam (Gasteiger et al., 2005) and net charge via the Henderson-Hasselbalch equation (Moore, 1985). To detect compositional bias in the protein sequences we used fLPS 2.0 (Harrison, 2021).

### Cloning of constructs

We synthesized complementary DNA (cDNA) with SuperScript™ II Reverse Transcriptase (Invitrogen) from total RNA extracted from *A. candida*-infected *A. thaliana* leaves at 8 dpi using an RNeasy kit (QIAGEN). We amplified the candidate sequences for the genes of interest with PCR using Phusion® High-Fidelity DNA polymerase (NEB) and primers designed for subsequent cloning (Table S4). We cloned the amplicons using the In-Fusion® cloning method (Takara Bio) into pET28b vectors for expression in *Escherichia coli* with two 6x Histidine tags (6xHis-tags) flanking the gene of interest for candidates C05, C06 and derivatives; and one 6xHis-tag at the C-terminus for the remaining candidates (C14 and derivatives, C15). We cloned candidates that had a putative secretion signal as predicted by SignalP version 5, namely C14 and C15, without the secretion-signal encoding sequence (Armenteros et al., 2019).

### Protein expression

We overexpressed the candidate proteins in the *E. coli* strain SHuffle® (NEB) at 30 °C (for candidates C14 and derivatives) and *E. coli* strain Rosetta™ DE3 (Merck) at 37 °C (for candidates C05, C15, C06 and derivatives). We induced expression at a 600 nm optical density (OD_600_) of 0.5-0.6 by adding isopropyl β-D-1-thiogalactopyranoside (IPTG) to a final concentration of 1 mm. We harvested 5 h post induction by collecting the pellets through centrifugation for 10 min at 8000 RCF, shock freezing and storing at −80 °C until further processing.

### Denaturing purification

We extracted the inclusion bodies from the cell pellets under denaturing conditions via sonication for 10 min (13 kHz in 0.6 s long pulses followed by 0.4 s of rest) in lysis buffer (100 mm sodium phosphate, 10 mm Tris, 7 m urea, pH 8) followed by shaking at room temperature for 1 h and centrifugation for 1 h at 10,000 RCF and 4 °C. We purified the proteins with HisTrap™ excel (GE Healthcare) in a one-step elution (elution buffer: 100 mm sodium phosphate, 10 mm Tris, 7 m urea, pH 4.5). We concentrated the elution by centrifugation in a VivaSpin® 20 column (Sartorius). We performed refolding by dialysis in a 3,500-5,000 molecular weight cutoff Float-a-lyzer® G2 device (Spectrum Labs). We gradually exchanged the denaturing elution buffer with the testing buffer (10 mm BisTris, pH 5.9) in four steps over a period of 48 h at 7 °C. We kept the final rebuffering solution in contact with the dialysis tube as a negative control for the antimicrobial activity tests. We determined the final protein concentration by Bradford assay using a Bovine Serum Albumin (BSA) standard (Thermo Scientific) and assessed protein purity by SDS-PAGE.

### Antimicrobial testing

We tested strains from a stock of plant-isolated bacteria collected during the *A. thaliana* sampling used to create the amplicon sequencing dataset mentioned above (Tables S3). The strain library comprises epi- and endophytic bacterial species isolated from either *Albugo*-infected or non-infected plants and includes the strains contained in the synthetic community (SynCom) mentioned in the next section. We applied a standard culturing approach to isolate strains from the *A. thaliana* samples. We used four different media for bacterial isolation (nutrient broth, Reasoner’s 2A agar, tryptic soy broth and King’s B agar) and cultivated at 22 °C. The strains used in this study were taxonomically assigned through similarity of their 16S region through blastn of the National Center for Biotechnology Information (NCBI) ribosomal RNA database. We grew overnight cultures from single colonies in nutrient broth medium at 22 °C. We adjusted protein molarity and added the rebuffered proteins at a volume ratio of 1:1 to the cell cultures diluted to a starting OD_600_ of 0.1 for a final testing volume of 100 μL. All tests were performed in at least three biological replicates in 96-well VWR transparent flat bottom plates. We used the TriStar2 plate reader (Berthold) with a program that measured OD_600_ every 15 min over 19 h with constant orbital medium strength shaking at room temperature (22 °C). We conducted bacterial community experiments in culture with two of the peptides (C06d and C06b). For these, we constructed a five-strain SynCom (I01, I03, I04, I13, I28; Table S1). We grew overnight cultures of the individual strains at 22 °C, adjusted their OD_600_ to 0.2 in fresh NB medium and mixed them at equal volume to form the SynCom, with a final OD_600_ of 0.02 for each strain in the mix. We combined 45 μL of the SynCom with 50 μL of buffer or protein solution adjusted to 1 μm final well concentration in 96-well VWR transparent flat bottom plates, followed by incubation at 22 °C and 180 rpm shaking for 24 h. We added 5 μL of a rifampicin-resistant *Pseudomonas syringae* pv. tomato DC3000 (Pst) culture (OD_600_ = 0.2) to a total of 100 μL per well and continued incubation for another 24 h. We created dilution lines of each well and dropped 10 μl of the dilutions on NB agar plates containing 50 μg mL^−1^ rifampicin. We counted Pst colonies after incubating the plates for 32 h h at 22 °C. The experiments were repeated three times with nine biological replicates per treatment and round. We performed the following treatment combinations: 1. SynCom plus buffer, 2. SynCom without I01 plus buffer, 3. SynCom plus peptide C06d, 4. SynCom plus peptide C06b.

For the *Aeromicrobium fastidiosum* (I01) interactions with other bacterial species from the SymCom, we examined one-to-one inhibition by a co-cultivation setup. We grew overnight cultures in NB medium and adjusted OD_600_ to 1.0. We spread a lawn of test strains on NB agar medium plates and aseptically created a hole with a sterile cork borer to dispense 100 μL of culture. We incubated the plates at 22 °C for three days to visualize the zone of inhibition.

### Synthetic microbial community experiment

For the SynCom experiment, we used *A. thaliana* plants of ecotype Ws-0. We sterilized the seeds with chlorine gas for 6 h (Lindsey et al., 2017). We further checked for bacterial and fungal seed-borne contaminants on NB and potato dextrose agar, respectively, by incubation for one week at 22 °C. After one week, seedlings were singularised and transferred to 1/2 Murashige and Skoog (MS) media in sterile 12-well plates (Greiner bio-one) and grown for three more weeks. The SynCom was assembled with core microbes from the wild *A. thaliana* population, consisting of 12 bacteria and three yeasts (Table S3). Members of the SynCom were grown and adjusted to an OD_600_ of 0.2 in 10 mm MgCl_2_ with 0.04% Silwet. They were mixed in equal parts and sprayed onto the 4-week-old seedlings. One week after initial SynCom inoculation, we sprayed the plants with rifampicin-resistant model pathogen Pst. Pst was grown in NB at 22 °C with shaking at 180 rpm and processed in the same way as the SynCom members. We sampled plants for amplicon sequencing at three time points, after SynCom spraying on day 0, on day 7 before inoculation with Pst and on day 10, third day post Pst infection. In total, three biological replicates were taken from each of the three treatments, whole SynCom, SynCom without *Aeromicrobium* (I01) and mock control without SynCom. DNA extractions were performed using a PowerSoil DNA Isolation Kit (MO BIO Laboratories Inc.). Amplicon libraries for sequencing were prepared using V5-V7 regions from the small prokaryotic subunit (16S rRNA gene) and ITS2 regions (internal transcribed spacer) for targeting bacterial and fungal amplicons, respectively (Agler et al., 2016). Sequencing was performed with Illumina MiSeq platform using V3 kit (600 cycles). Finally, we processed and analyzed the raw sequencing reads using QIIME2, with which we calculated Pst relative abundances and performed principal coordinate analysis based on Euclidean distances of the rarefied amplicon sequencing variant (ASV) frequencies to measure sample to sample variation (Bolyen et al., 2019).

## Data availability

All data discussed in this paper as well as the code to reproduce the analyses and figures are found at https://doi.org/10.5281/zenodo.6325163. The mass spectrometry proteomics data have been deposited to the ProteomeXchange Consortium via the PRIDE partner repository with the dataset identifier PXD031981 (Perez-Riverol et al., 2021).

## Acknowledgements

We would like to acknowledge support from the graduate school GRK 1708 “Molecular principles of bacterial survival strategies” and the Open Access Publishing Fund of the University of Tübingen. We would like to thank Libera Lo Presti for her comments and suggestions on the manuscript. Furthermore, we would like to thank Maryam Mahmoudi for providing the raw OTU tables from the amplicon sequencing, Samuel Kroll, Paul Runge and Jonas Ruhe for isolating and sequencing some of the tested bacterial strains and Sophia Häußler for her help during the SynCom experiments.

## Supplementary figures

**Figure S1.**
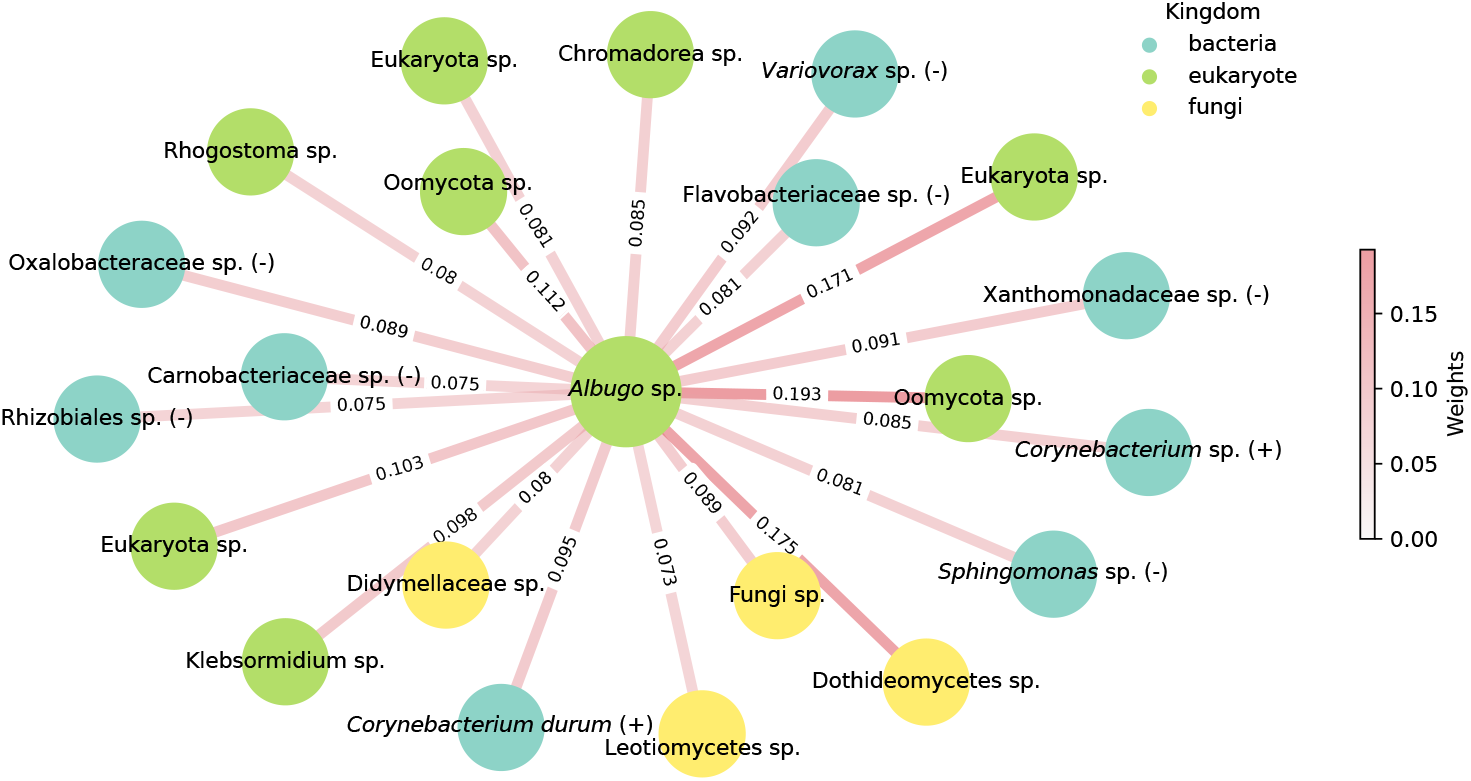
Inferred positive interactions for the *Albugo* sp. operational taxonomic unit on the *Arabidopsis thaliana* phyllosphere amplicon dataset. Color of the edges represents the strength of the correlation and color of the nodes represents phylogenetic kingdom. Unless explicitly stated, taxonomy at the species level could not be resolved with confidence (bootstrap < 90).

**Figure S2.**
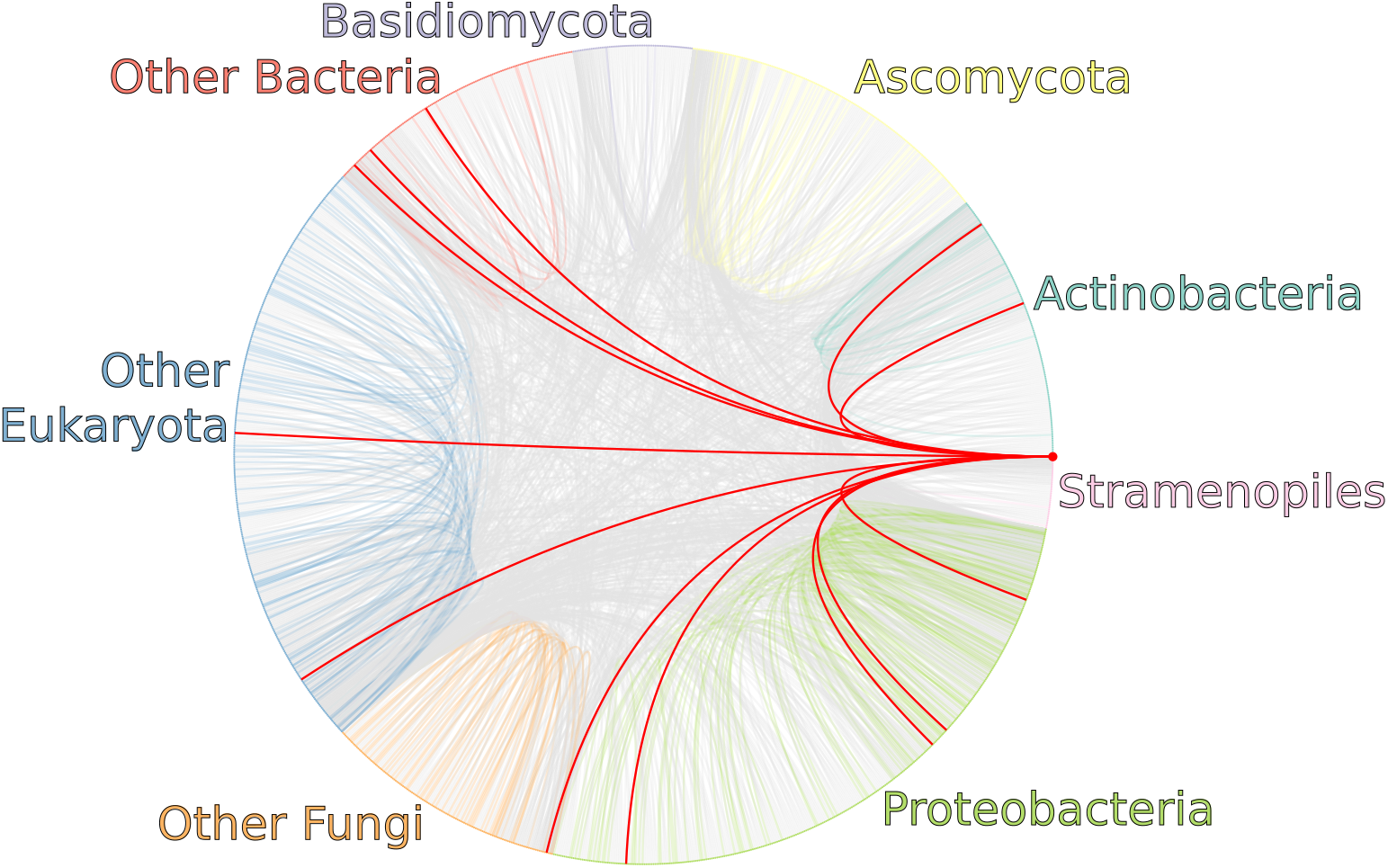
Overview of negative correlation network inferred by FlashWeave on the *Arabidopsis thaliana* phyllosphere amplicon dataset. Operational taxonomic unit (OTU) nodes are grouped based on phylogenetic similarity and ordered by edge count. Highlighted are the *Albugo* sp. OTU and its connecting edges. Intergroup edges are colored gray while intragroup edges have the same color as the corresponding nodes.

**Figure S3.**
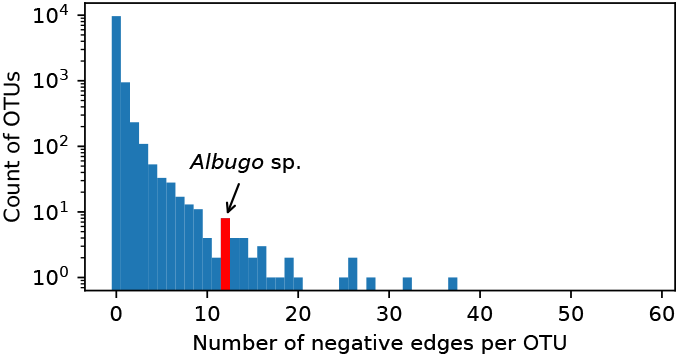
Histogram of the number of negative correlations as predicted by FlashWeave on the *Arabidopsis thaliana* phyllosphere amplicon dataset. Highlighted is the location of the *Albugo* sp. operational taxonomic unit (OTU).

**Figure S4.**
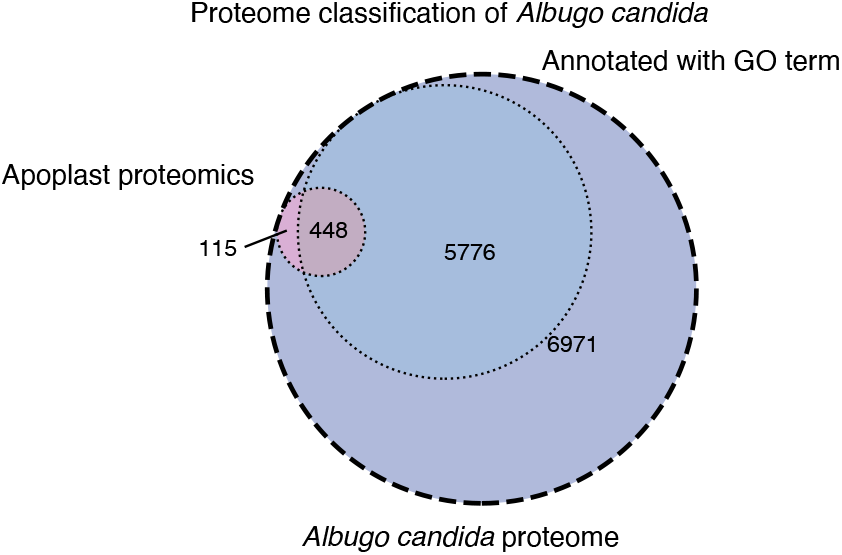
Venn diagram of annotated proteins in *Albugo candida*’s predicted proteome. Proteins found in the apopolast are pink and proteins annotated with at least a gene ontology (GO) term are in blue.

**Figure S5.**
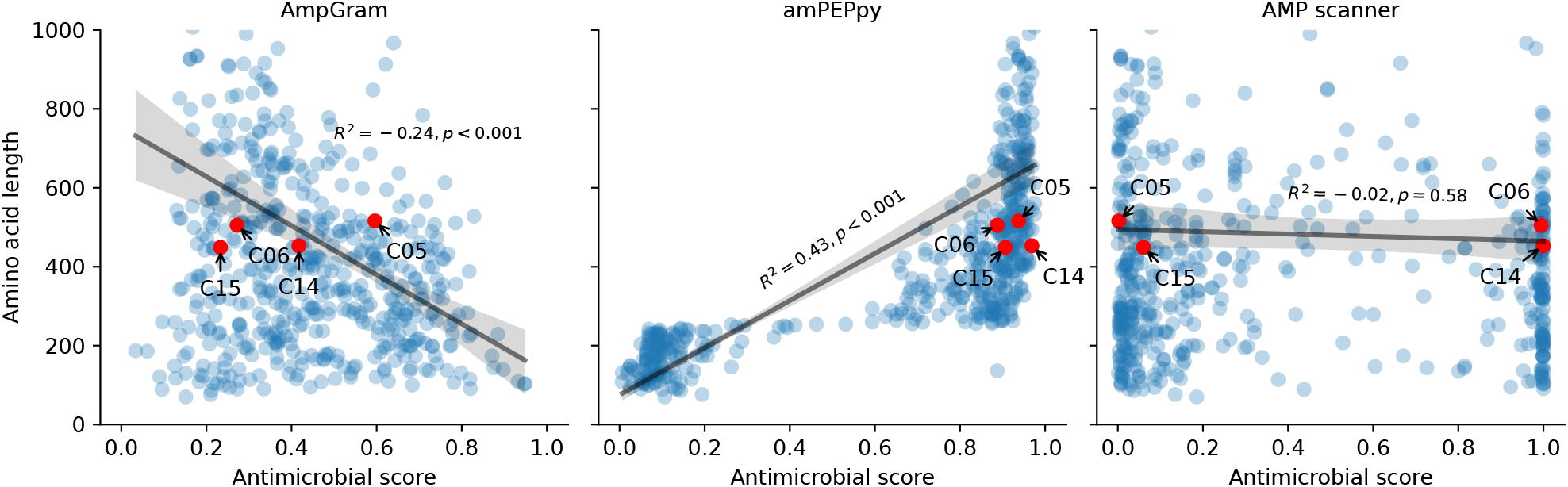
Scatter plot comparing the three machine learning methods for antimicrobial prediction to protein length in the proteins found in the apoplast for *Albugo candida*. Highlighted are the selected candidate proteins. The y-axis is displayed up to a limit of 1000 amino acids. The black line represents the best least-squares fit for linear regression. Shaded is the 95% confidence interval for each regression. Depicted are the R-squared parameter and the *p* value for Wald test with t-distribution using as null hypothesis a zero-slope regression. Note the significant correlation of AmpGram and amPEPpy to protein length.

**Figure S6.**
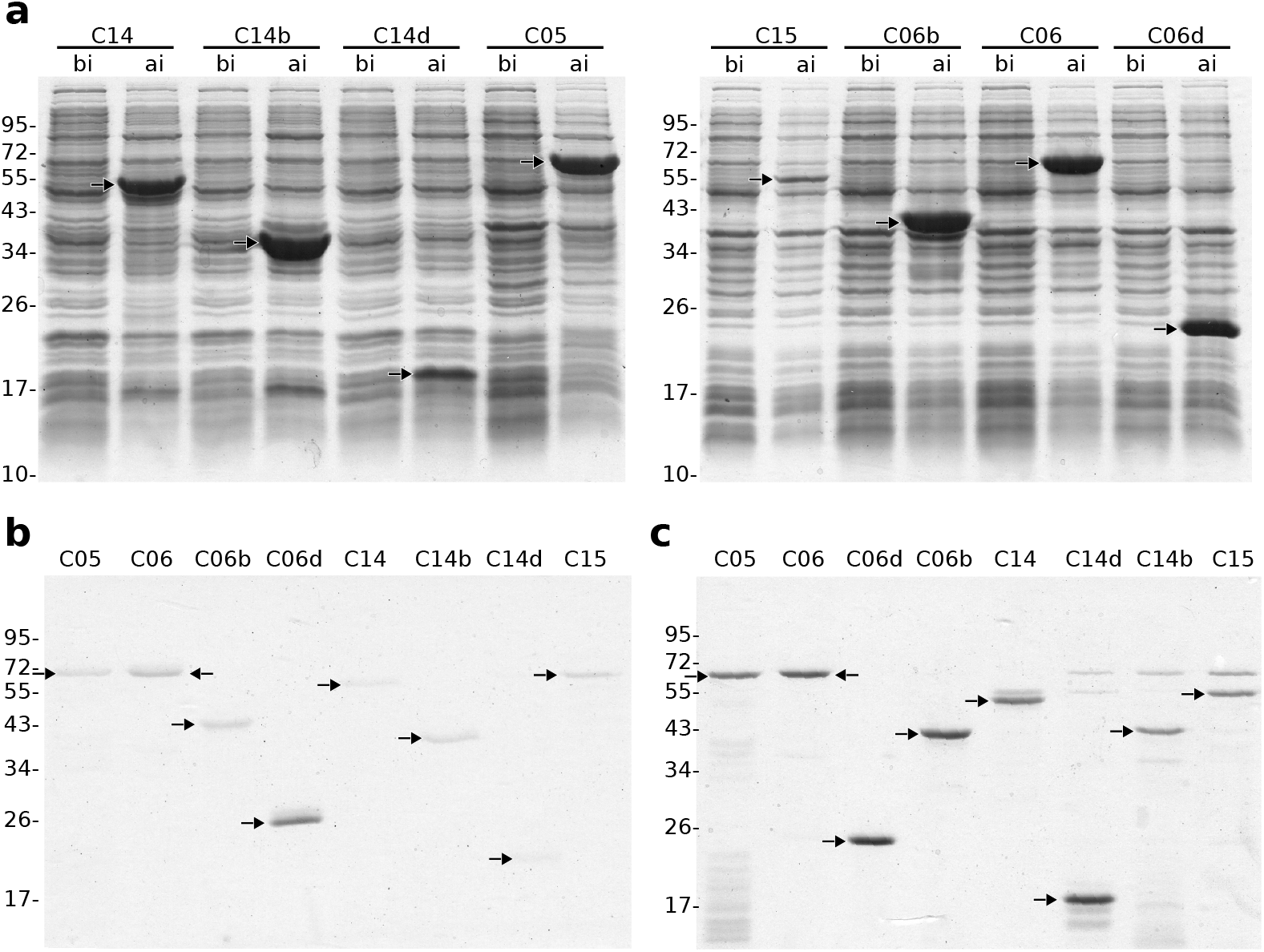
Expression and purification analysis of the protein candidates by SDS-PAGE and Coomassie blue staining. Scale on the left side indicates molecular weight in kDa. Black arrows highlight relevant bands. (**a**) Comparison of protein expression before (bi) and after (ai) IPTG induction of *Escherichia coli*. (**b**) Analysis of candidate protein elution fractions after purification in the denaturing buffer. (**c**) Analysis of protein candidates after concentration and rebuffering in the testing buffer.

**Figure S7.**
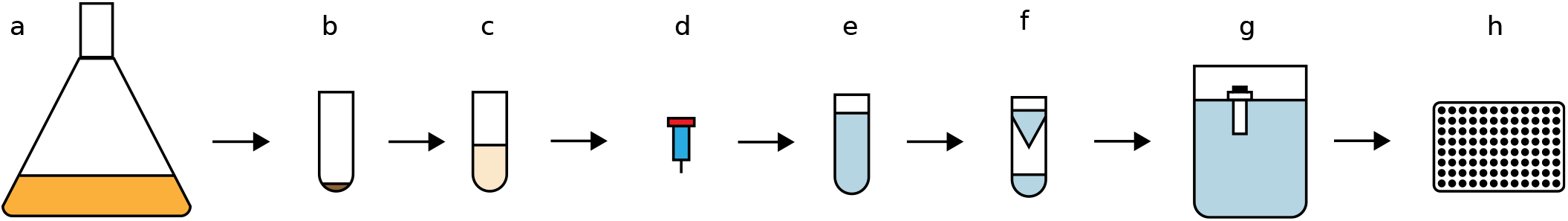
Workflow of the denaturing purification process and subsequent in vitro activity assays. (**a**) Overexpression of the candidate proteins in *Escherichia coli*. (**b**)Freezing of harvested pellets and storage at −80 ºC. (**c**) Resuspension in denaturing buffer and sonication to dissolve inclusion bodies. (**d**) Denaturing purification using a HisTrap™. (**e**) One-step elution using a low pH buffer. (**f**) Concentration of denatured protein in a VivaSpin® 20 column. (**g**) Dialysis of protein over two days in increasing concentrations of refolding buffer. (**h**) Antimicrobial assays on bacterial strains.

**Figure S8.**
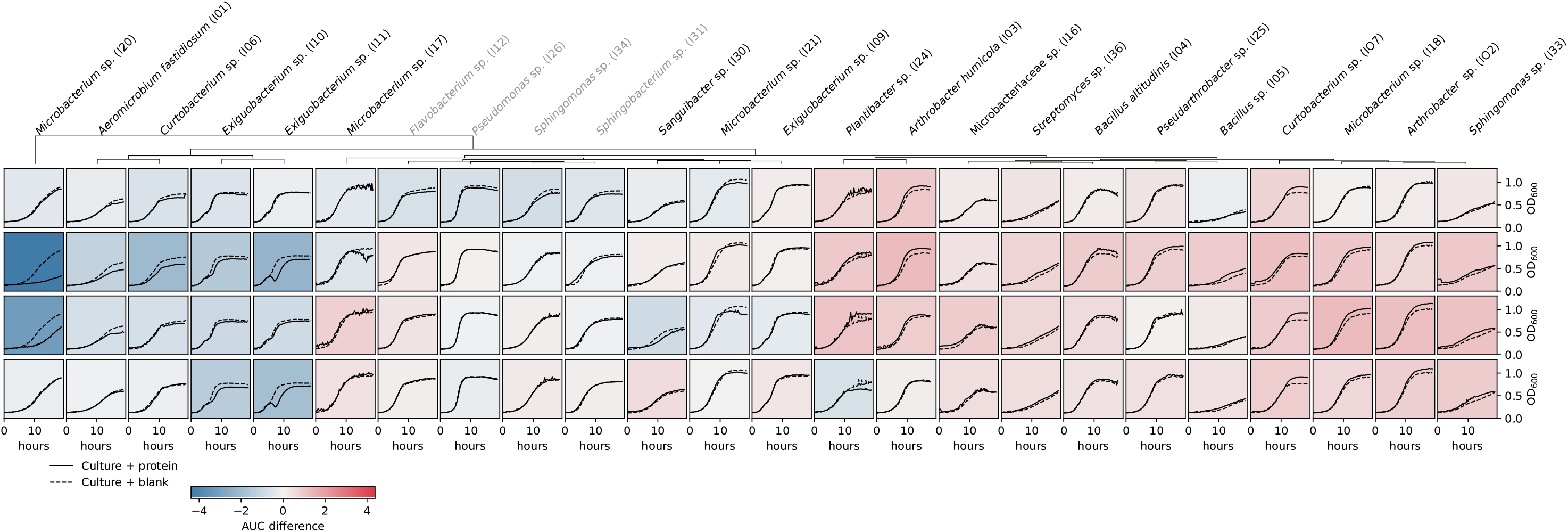
Growth curves for candidates C05, C06, C14 and C15 (from top to bottom row) at a concentration of 0.75 mM compared to blank (dashed lines). Background color represents inhibition (blue) or promotion (red) of growth based on the difference in the area under the curves of the means of three biological replicates. Names in gray represent gram-negative bacterial strains, while the rest are gram-positive.

**Figure S9.**
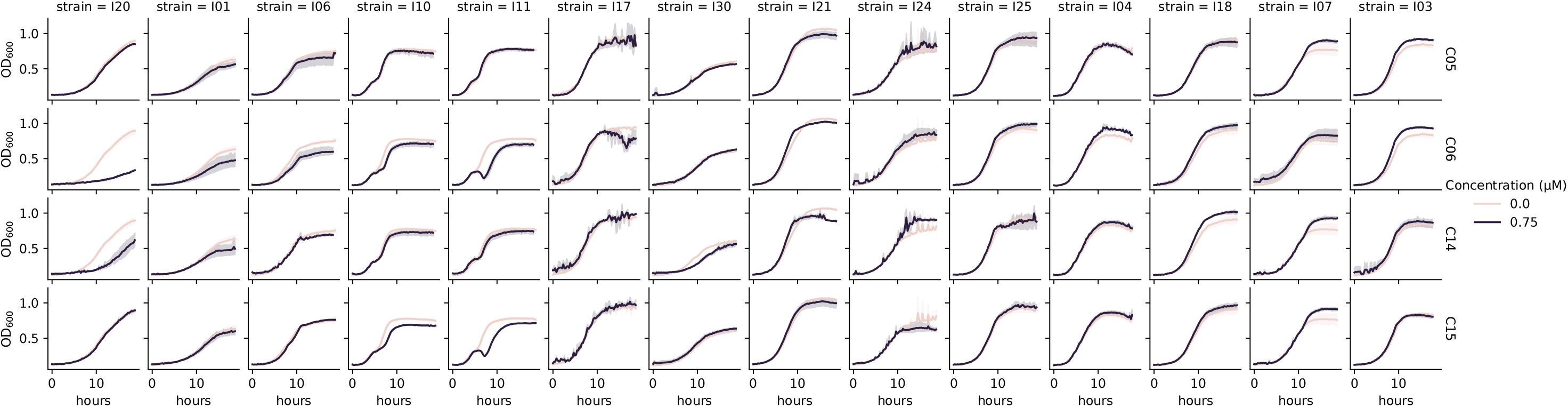
Growth curves for candidates C06, C06, C14 and C15 (from top to bottom row) in purple at a concentration per well of 0.75 μM compared to blank (beige). Background color represents inhibition (blue) or promotion (red) of growth based on the difference in the area under the curves of the means of three biological replicates. Confidence intervals of 95% from at least three biological replicates shown as the colored area around the mean.

**Figure S10.**
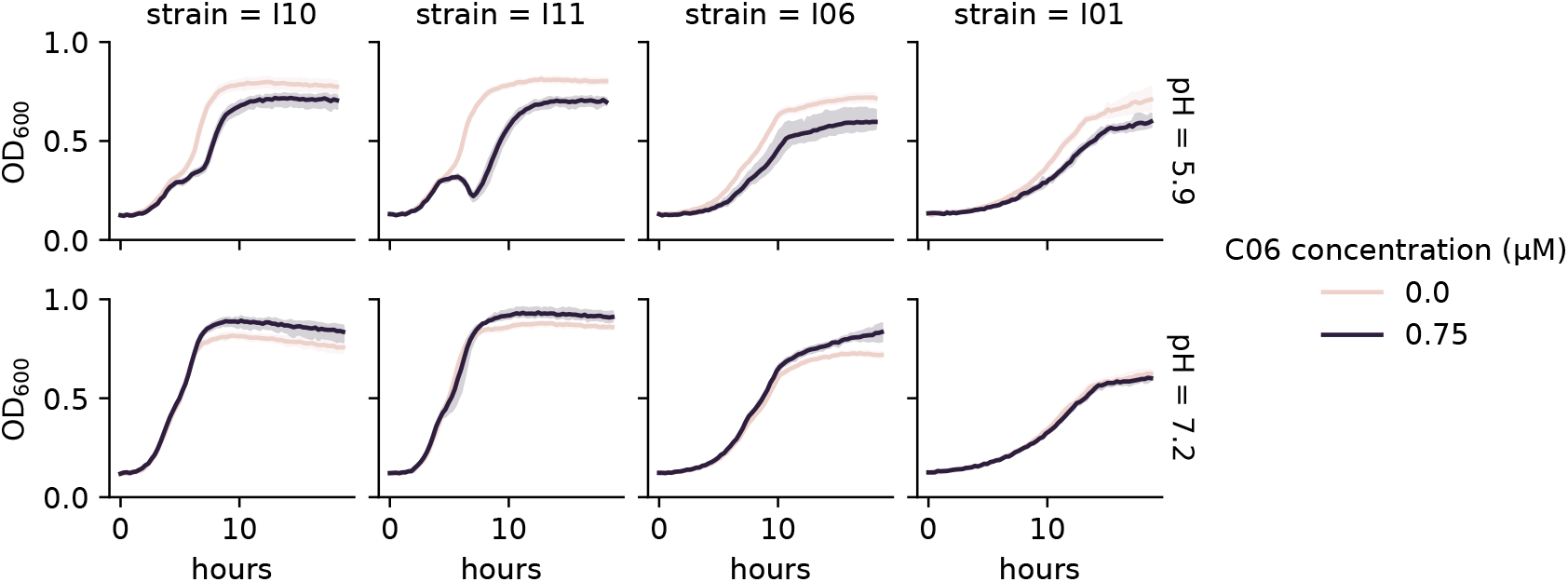
Comparison of the antimicrobial activity of C06 at pH 5.9 and 7.2 at molarity 0.75 μM. Tested strains included *Exiguobacterium* I10 and I11, *Curtobacterium* I06 and *Aeromicrobium fastidiosum* I01. Confidence intervals of 95% from at least six biological replicates shown as the colored area around the mean. Blank treatments are represented as beige lines and protein treatments in purple.

**Figure S11.**
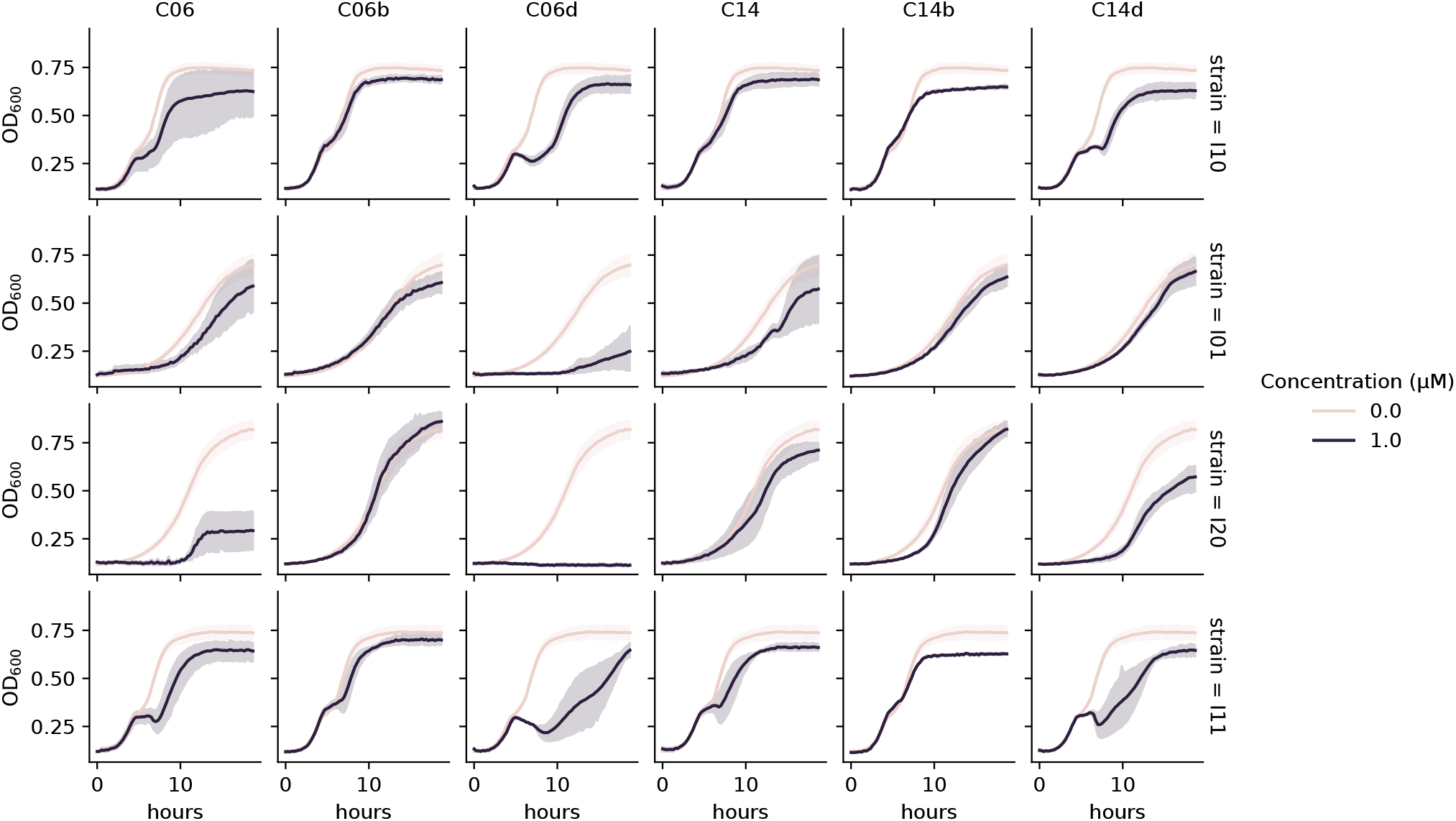
Inhibition curves of C06, C14 and their domains at a concentration of 1 μM. The tested strains are represented in different rows as follows (from top to bottom): *Exiguobacterium* sp. I10, *Aeromicrobium fastidiosum* I01, *Microbacterium* sp. I20 and *Exiguobacterium* sp. I11. Confidence intervals of 95% from at least six biological replicates shown as the colored area around the mean. Blank treatments are represented as beige lines and protein treatments in purple.

**Figure S12.**
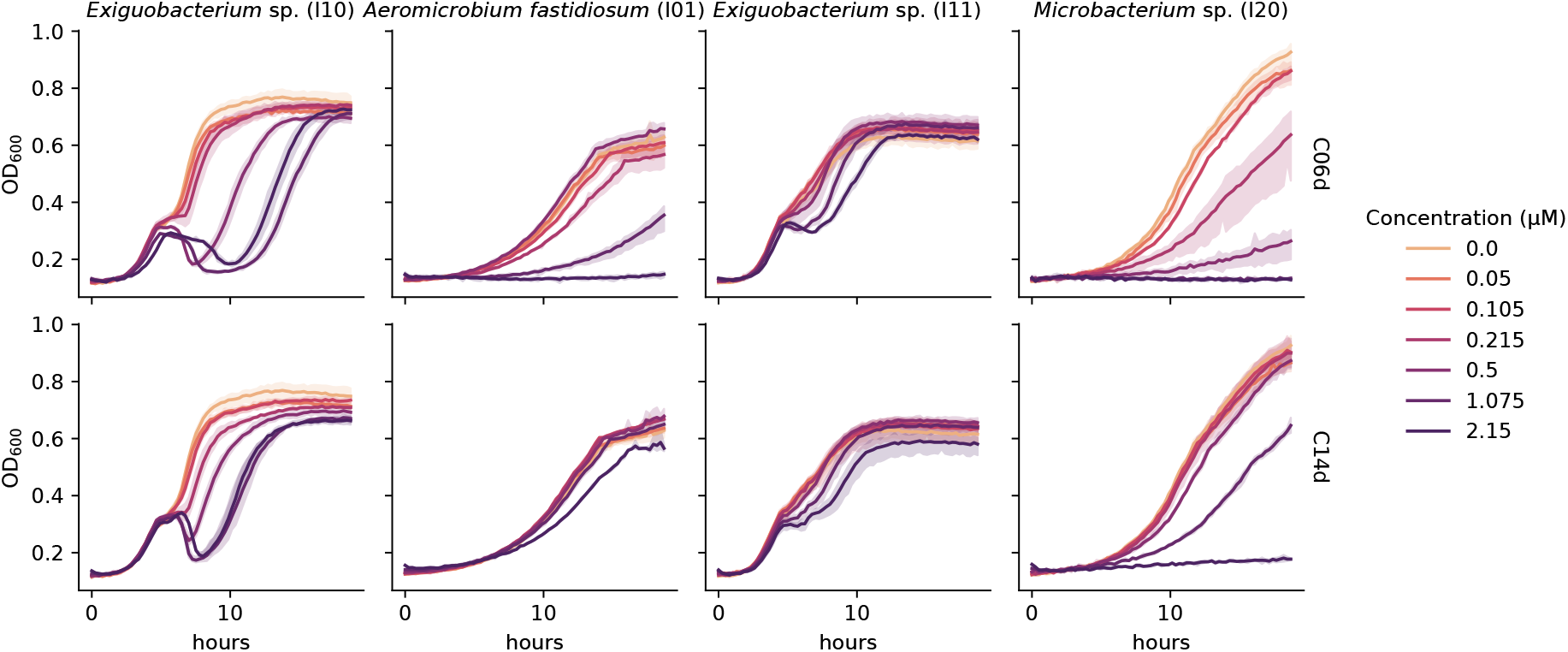
Inhibition curves of six different concentrations of C06d and C14d domains (first and second row, respectively) towards four sensitive strains. Confidence intervals of 95% from three biological replicates shown as the area around the mean. The color gradient represents the concentration of proteins while beige is the blank.

**Figure S13.**
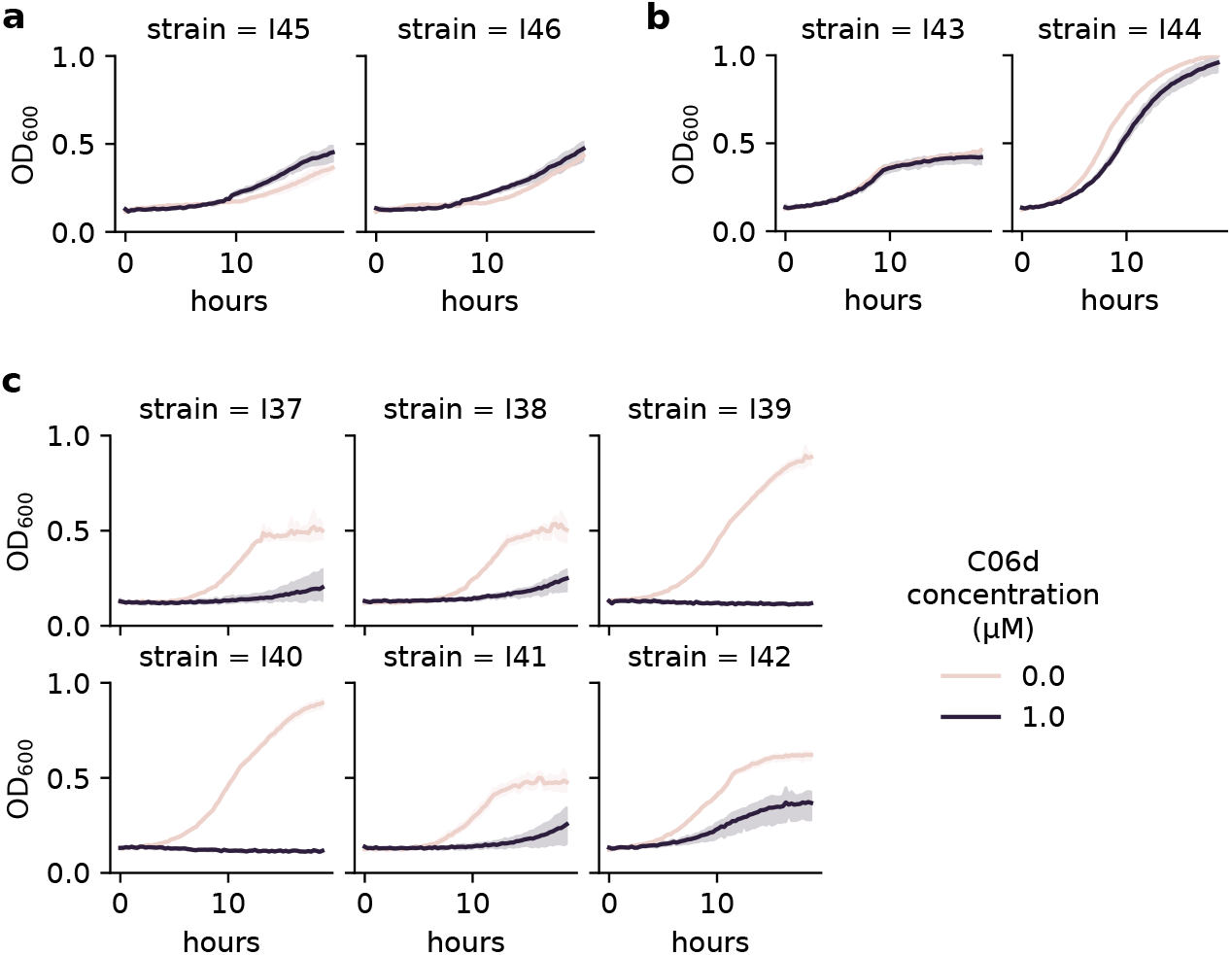
Inhibition curves of C06d on *Clavibacter* and *Rhodococcus* strains at a concentration of 1 μM (purple) and 0 μM (beige). Confidence intervals of 95% from three biological replicates shown as the shaded area around the mean. (**a**) Growth curves of isolates from *Clavibacter michiganensis* subsp. *tessellarius*. (**b**) Growth curves of isolates from *Rhodococcus fascians*. (c) Growth curves of isolates from *C. michiganensis* subsp. *capsici*.

**Figure S14.**
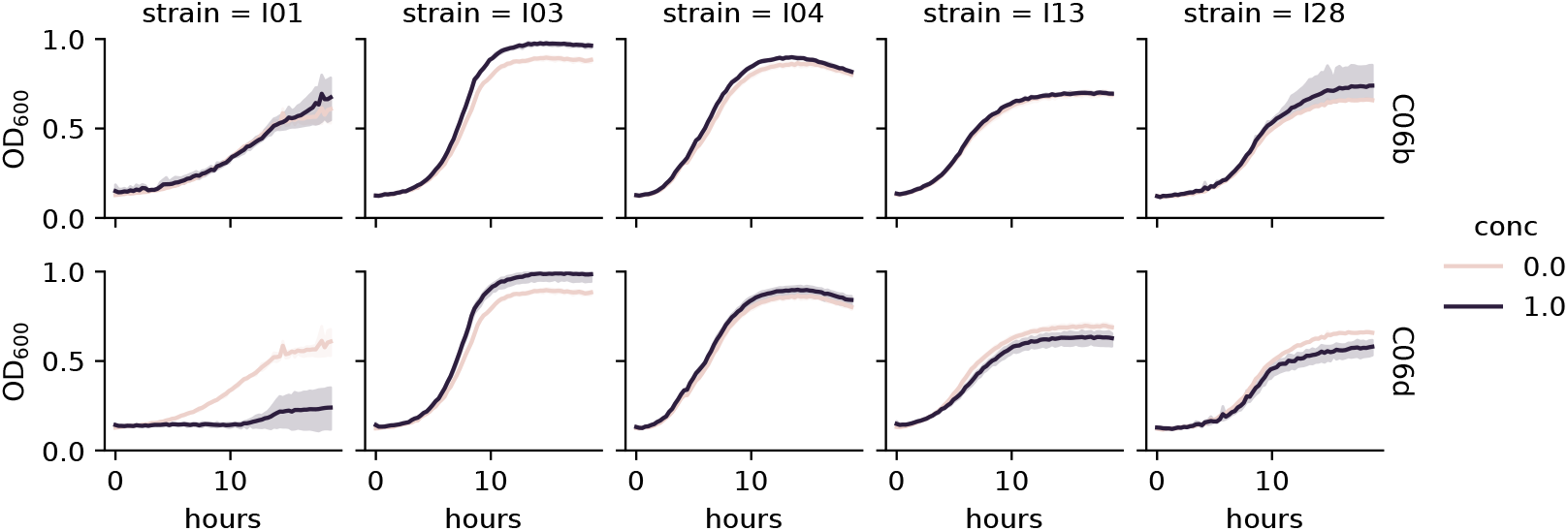
Growth curves from single-strain antimicrobial activity assays with the individual strains from the five-strain bacterial community and added C06b (first row) and C06d (second row) at a concentration of 1 μM (purple) and 0 μM (beige).

**Figure S15.**
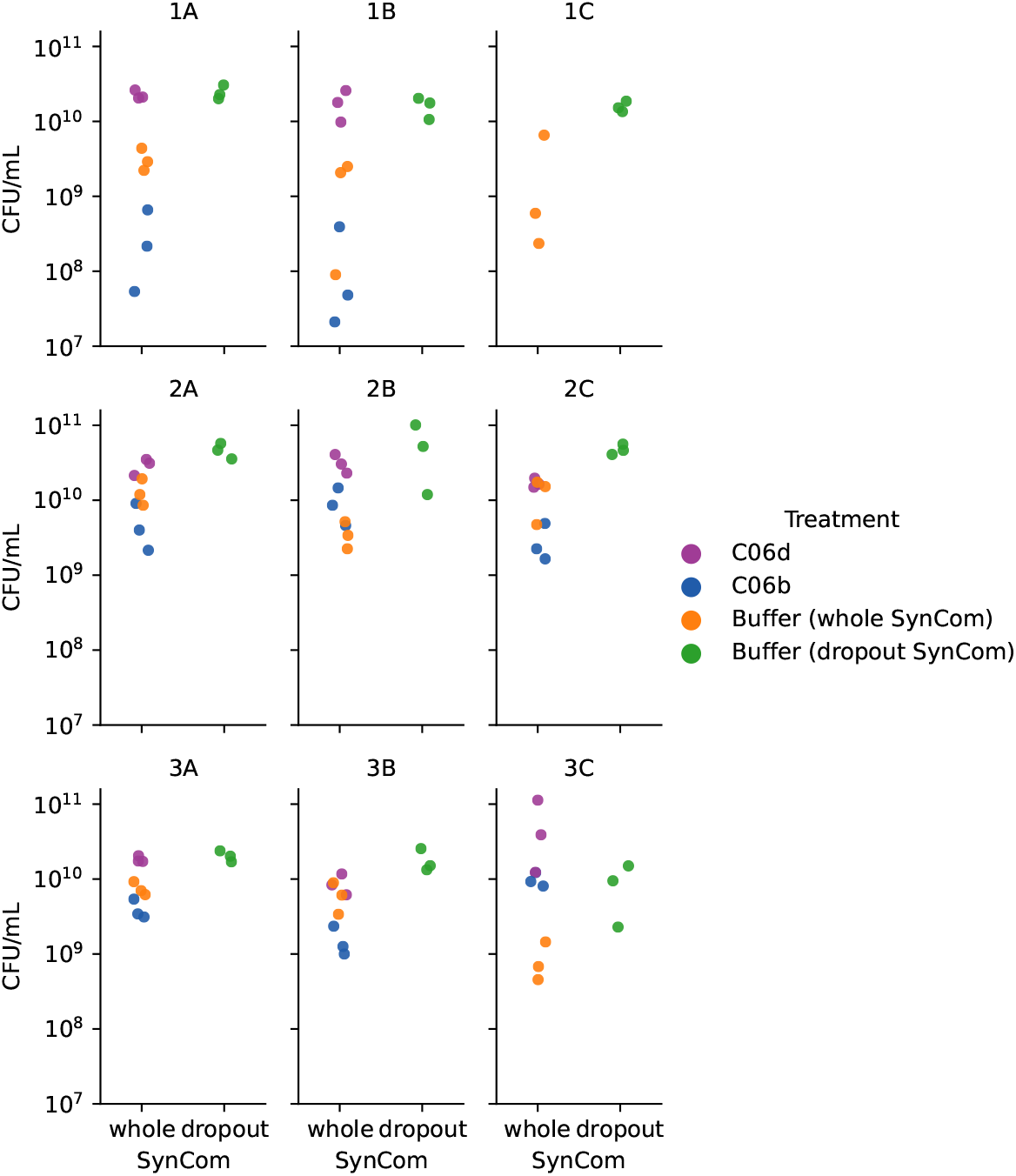
CFU counts per replicate of *Pseudomonas syringae* pv. Tomato DC3000 (Pst) in five-strain bacterial synthetic community (SynCom) experiments in liquid culture. Effects of C06d (purple) and C06b (blue) candidates at a concentration of 1 μM on whole SynCom treatments are displayed. Effects of the buffer (BisTris, pH 5.9) on the whole community are shown in orange and on the dropout community in green. The x-axis displays community status (‘whole’ for whole five-strain community, ‘dropout’ for community without *Aeromicrobium fastidiosum* I01). Rows represent time points of replicates, and columns, different biological replicates for the same time point.

**Figure S16.**
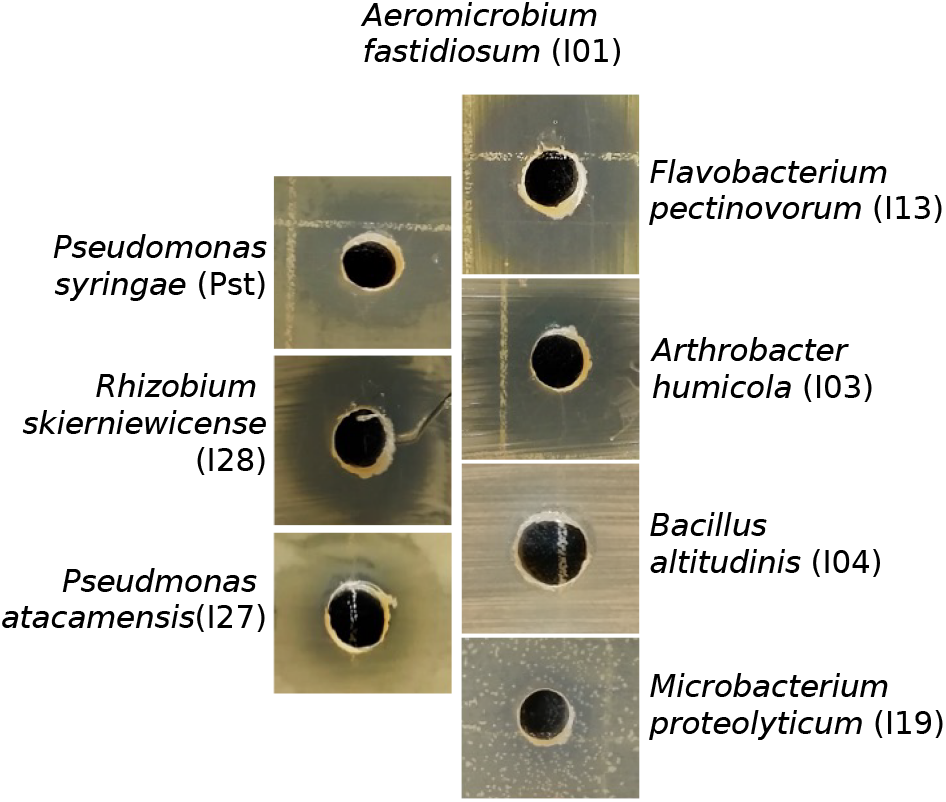
Pairwise inhibition rings of *Aeromicrobium fastidiosum* (I01) against other strains in the synthetic bacterial community.

**Figure S17.**
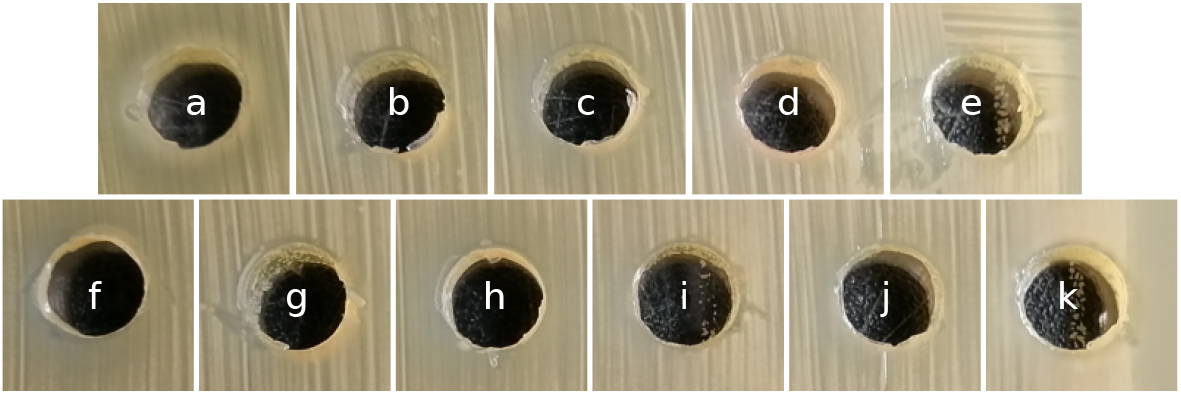
*Aeromicrobium fastidiosum* I01 lawn for interaction testing with bacterial strains in the synthetic bacterial community. (**a**): *Pseudomonas atacamensis* (I27), (**b**): *Flavobacterium pectinovorum* (I13), (**c**): *Rhizobium skierniewicense* (I28), (**d**): *Microbacterium proteolyticum* (I19), (**e**): *Arhtrobacter humicola* (53), (**f**): *Bacillus altitudinis* (I04), (**g**): *Frigoribacterium faeni* (I14), (**h**): *Sphingomonas faeni* (I32), (**i**): *Nocardioides cavernae* (I22), (**j**): *Paenibacillus amylolyticus* (I23), (**k**): *Methylobacterium goesingense* (I15).

**Figure S18.**
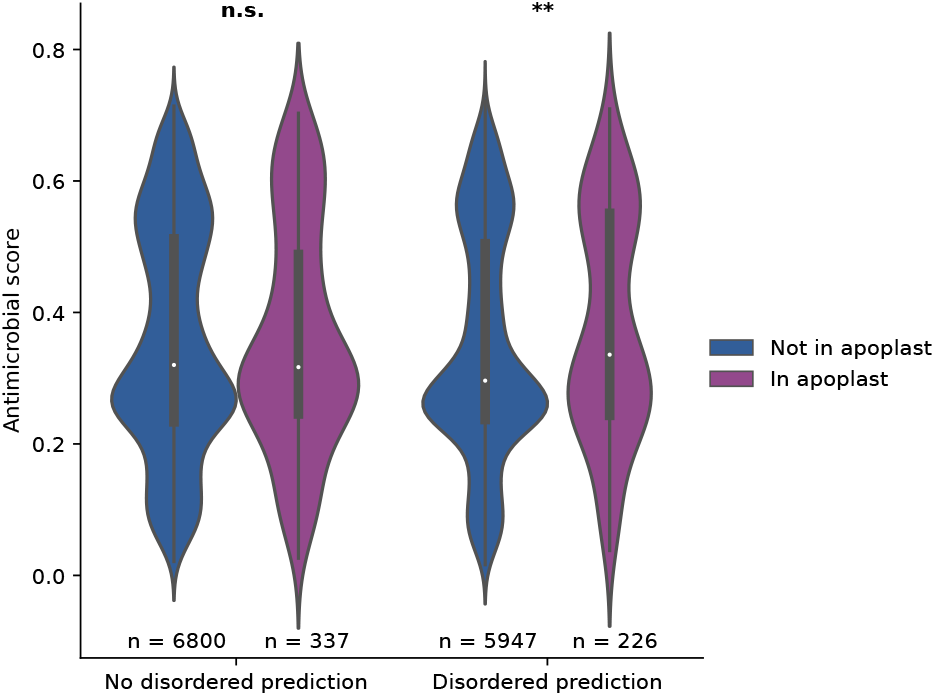
Antimicrobial prediction of apoplastic and non-apoplastic proteins grouped by the presence or absence of intrinsically disordered regions *Albugo candida*. Significance was tested using Holm-corrected two-tailed Mann-Whitney U tests (*p* value = 0.008 for disordered prediction). *n*.*s*.: *not significant, **: p value < 0*.*01*.

## Supplementary tables

**Table S1.**
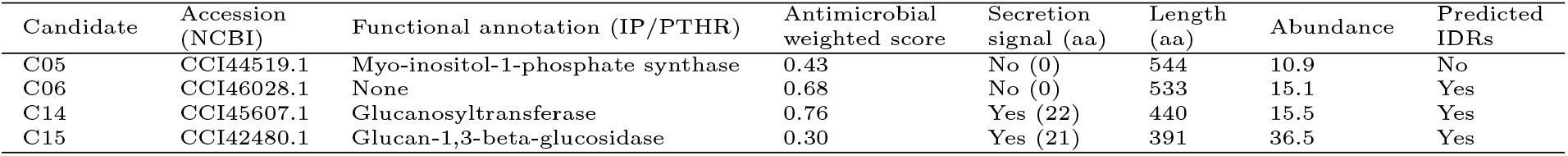
Apoplastic protein candidates from *Albugo candida* selected for antimicrobial testing. Abundance in the apoplast is represented as the ratio of the intensity peptide values comparing infected and uninfected *Arabidopsis thaliana* of the three replicates after normalization. *NCBI: National Center for Biotechnology Information, IP: InterPro, PTHR: Panther, aa: amino acid, IDR: intrinsically disordered region*.

**Table S2.** Net charge at pH 5.9 and 7.2, theoretical isoelectric point (pI) and molecular weight in kilodalton (kDa) of expressed candidate proteins and domains.

| Candidate | Theoretical pI | Net charge (pH 5.9) | Net charge (pH 7.2) | Molecular weight (kDa) |
| --- | --- | --- | --- | --- |
| C05 | 6.27 | 5.90 | —7.92 | 60.2 |
| C06 | 6.28 | 8.05 | —9.96 | 57.1 |
| C06d | 6.86 | 13.55 | —1.27 | 21.2 |
| C06b | 6.05 | 1.76 | —7.92 | 39.4 |
| C14 | 6.23 | 2.85 | —5.27 | 48.9 |
| C14d | 8.79 | 8.03 | 4.00 | 15.1 |
| C14b | 5.60 | —2.82 | —9.91 | 35.0 |
| C15 | 5.89 | —0.36 | —12.77 | 49.0 |

**Table S3.**
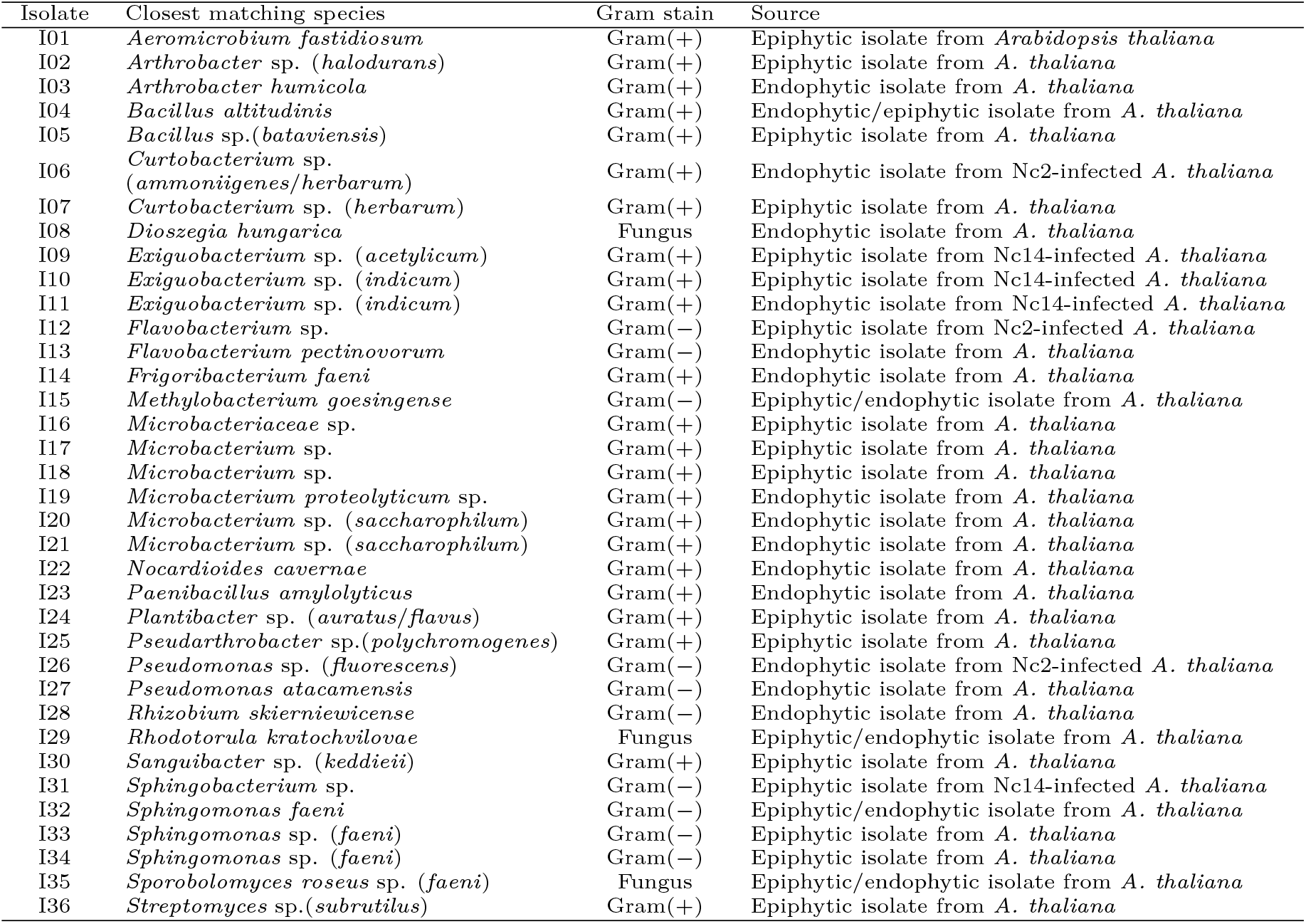
Strain collection of plant-isolated microbes from environmental sampling of Arabidopsis thaliana. It includes strains from the *A. thaliana* core synthetic community (SynCom); I03: GCF_022803015.1, I04: GCF_022803025.1, I13: GCF_022802975.1, I14: GCF_022805035.1, I15: GCF_022803065.1, I19: GCF_022803055.1, I27: GCF_022802935.1, I28: GCF_022802995.1, I29: GCA_022817825.1, I35: GCA_022817875.1. Taxonomic assignment was based on blastn matches on the 16S database from the National Center for Biotechnology (NCBI), as of February 2022. Taxonomy assignment of SynCom strains was based on their whole genome sequencing data. In parentheses, the closest matching species name is depicted. Fungal sequences were blasted against the non-redundant nucleotide database from NCBI. *Nc2: Albugo candida strain Nc2, Nc14: Albugo laibachii strain Nc14*.

**Table S4.**
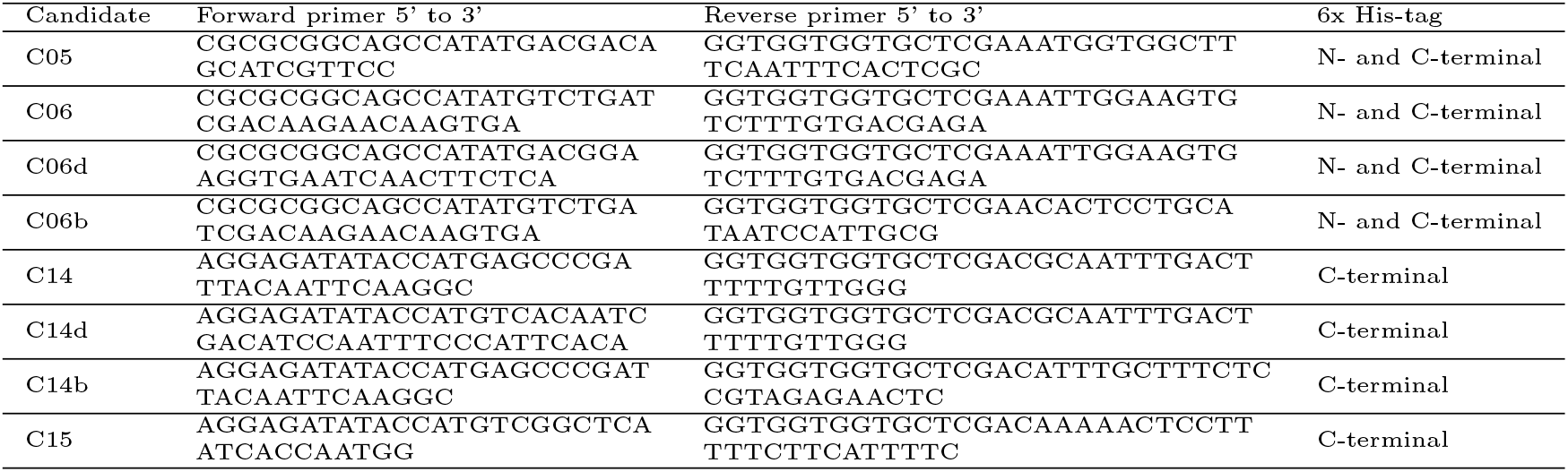
Primer sequences for amplification and subsequent cloning of candidate sequences from *Albugo candida* into vector pET28b.

**Table S5.** Significant compositional bias of tested protein candidates. Significance given as the negative logarithm of the binomial *p* values.

| Candidates | Residues | Significance |
| --- | --- | --- |
| C05 | N | 7.0 |
| C06 | A, Q | 15.8, 12.4 |
| C06b | Q, A | 11.7, 9.3 |
| C06d | H, A | 13.5, 9.0 |
| C14 | S, C | 10.8, 7.2 |
| C14d | S | 14.8 |
| C15 | S | 10.3 |

**Table S6.**
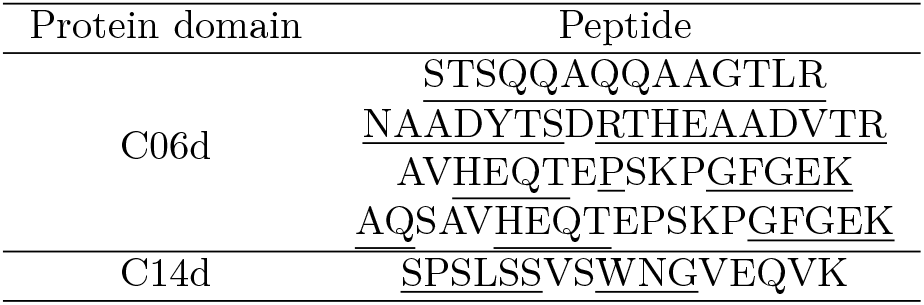
Peptide coverage of the N-terminal disordered domains from candidate proteins in the proteomics analysis. Underlined are residues predicted to be disordered.

**Table S7.**
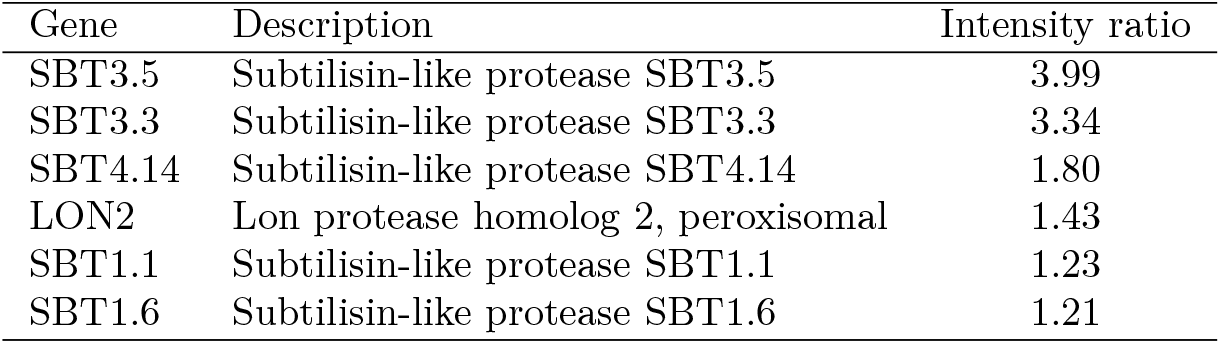
Proteases from *Arabidopsis thaliana* enriched in the *Albugo candida*-infected treatment.

**Table S8.** Plant-isolated putative pathogenic gram-positive microbes from environmental sampling of Arabidopsis thaliana employed in the testing of *Albugo candida* peptide C06d. Taxonomic assignment based on results of blastn on the 16S database from the National Center for Biotechnology, as of March 2022.

| Isolates | Closest matching species | Pathogenic phenotype | Reference |
| --- | --- | --- | --- |
| I37,I38,I39,<br>I40,I41,I42 | *Clavibacter michiganensis*<br>ssp. *capsici* | Pepper pathogen causing<br>bacterial canker disease | Type strain PF008 (Oh et al., 2016) |
| I43,I44 | *Clavibacter michiganensis*<br>ssp. *tessellarius* | Wheat pathogen causing<br>leaf freckles and spots | Strain 78181/ DSM 20741/ATCC 33566<br>(Carlson and Vidaver, 1982; Li and Yuan, 2017) |
| I45,I46 | *Rhodococcus fascians* | Broad host range pathogen<br>causing leafy gall disease | Type strain DSM 20669/CF17<br>(Dhaouadi et al., 2020; Hjerde et al., 2013) |

